# Sex and regional effects of *Bacteroides* in the gut

**DOI:** 10.1101/2025.08.27.672693

**Authors:** Rebecca A. Valls, Kaitlyn E. Barrack, Sarvesh V. Surve, James B. Bliska, George A. O’Toole

**Author notes:** Address correspondence to: George A. O’Toole, Dept. of Microbiology & Immunology, Rm 202 Remsen Building, 66 College Street, Geisel School of Medicine at Dartmouth Hanover, NH 03755.

## Abstract

*Bacteroides* spp. is a key immune-programming microbe in healthy individuals – these bacteria have been shown to be reduced in abundance across a variety of disease states. Our study investigated the systemic and region-specific responses to *Bacteroides* colonization in the gut, including sex-related differences, in mice. Utilizing C57BL/6 mice, we administered *Bacteroides* to conventional, antibiotic-treated mice, then assessed this microbe’s influence on the gut microbiota composition and inflammatory responses following an airway lipopolysaccharide challenge to assess effects on the gut-lung axis. We found that *Bacteroides* successfully colonizes the intestinal tract of antibiotic-treated mice, particularly the colon lumen of the large intestine as evidenced by 16S rRNA amplicon gene sequencing and culturing. Differential gene expression analysis using NanoString technology revealed significant immune response variations across the gut regions, with notable differences in adaptive immune response genes. A striking sex-dependent outcome was noted in the regulation of *atg12* in the cecum, potentially enhancing autophagic function, particularly in female mice. Additionally, *Bacteroides* intestinal colonization was associated with altered expression of macrophage markers such as *cd163*, *cd84*, and *ms4a4a*, which may reflect shifts in the macrophage profile within the cecum. These findings pave the way for novel therapeutic approaches that leverage microbial impacts on gut and systemic health, offering a deeper understanding of *Bacteroides*’ role in human health and disease. Our study highlights the necessity for further research to elucidate the intricate relationships between gut microbiota, host immunity, biological sex and their interplay.

**Importance:** This research marks an investigation into how specific microbiota, like *Bacteroides*, regulate host responses across different gut regions to influence systemic health. By dissecting the impact of *Bacteroides* across multiple regions of the intestinal tract, this study offers new insights into the localized and whole-body effects of this important immune-programming microbe. Such an understanding is crucial as it helps in unraveling the complex interplay between gut microbes and the host’s immune system. This research helps bridge the gap between local intestinal ecology and overall systemic health, addresses important questions relevant to the gut-lung axis, and helps pave the way for innovative therapies.

## Introduction

The gut microbiota’s role in influencing host immune responses and overall health is increasingly evident. Our previous studies highlighted the key role of *Bacteroides* spp., particularly through its secreted metabolite propionate, in reducing inflammation in intestinal epithelial cells^1^. When introduced into a mouse model, *Bacteroides* spp. not only increased stool propionate but also reduced systemic inflammatory markers following pulmonary lipopolysaccharide (LPS) challenge; this reduced airway inflammation is propionate-dependent. While propionate could be acting directly on airway immune cell function, we also explored the possibility that this short chain fatty acid (SCFA) might be acting locally in the gut. Additionally, the presence of *Bacteroides* appears to suppress the abundance of potentially harmful bacteria like *Escherichia-Shigella*, indicating that this anaerobe might impact intestinal microbial ecology.

The gastrointestinal tract’s distinct regions each present unique ecological niches, impacting the composition and function of the resident microbiota. In the small intestine, rapid content transit, a rich landscape of antimicrobial peptides, and a complex immune environment combine to influence microbial composition and activity. Shifts in environmental conditions are pronounced for transition to the cecum, where lower pH and a highly anaerobic environment prevail. The colon, traversing ascending to descending sections, exhibits a slow transit, encouraging extensive microbial fermentation and digestion processes. These variations in the small intestine, cecum, and colon are crucial for understanding the intricate dynamics of microbiota-host interactions^2,3^.

*Bacteroides spp.* serve as pivotal members of the gut microbiota, exhibiting a complex role in host health. On one hand, this microbe provides colonization resistance against pathogens through immune modulation and competition for nutrients. On the other, it can facilitate infection by cross-feeding pathogens and aiding their niche localization. This functional dynamic highlights the context-dependent potential of *Bacteroides* spp. to either protect the host or contribute to disease progression^4,5^. The beneficial functions of *Bacteroides spp.* include the degradation of complex polysaccharides, T-cell activation, development of the gut associated lymphatic tissue, maturation of the immune system (which may prevent allergy and asthma), stimulation of Paneth cells to produce antimicrobial molecules, as well as potentially playing a role in preventing obesity^6^. Notably, depletion of *Bacteroides* spp. represents a hallmark of gut microbiome dysbiosis in both children and adults with cystic fibrosis (CF) ^7,8^. Similar reductions in *Bacteroides* spp. have been linked to prediabetes and insulin resistance in young and middle-aged adults^9^. A metanalysis revealed disequilibrium and depletion of *Bacteroides* in individuals with Crohn’s disease and ulcerative colitis^10,11^. Thus, the impact of the loss of *Bacteroides* may extend to multiple diseases.

The complex relationship between the microbiota and host health is key to understanding the pathophysiology of various diseases, emphasizing the significance of localized microbial impacts in the intestine with potential systemic effects. Here we focus on unraveling how *Bacteroides* colonization affects host immune responses across different intestinal regions and the associated systemic health implications, considering the role of sex. To address these questions, we treated C57BL/6 conventional mice with antibiotics to suppress their native intestinal microbiota, then administered either PBS or a *Bacteroides* cocktail in PBS via oral gavage. We then dissected the gastrointestinal tract into segments: the small intestine, cecum, proximal colon, and distal colon. From each segment, we collected both luminal and mucosal samples. These samples were analyzed to determine their microbial compositions and immune gene expression profile. We applied mixed-effects models to evaluate the impact of different variables, including treatment, biogeography, and sex. This approach allowed us to gain an in-depth understanding of the influence of *Bacteroides* on the various regions of the murine gut in terms of both the microbial and host responses and revealed potential sex-specific responses to these microbes.

## Results

### Impact of *Bacteroides* supplementation on gut microbiota is sex-dependent

We aimed to examine the influence of *Bacteroides* supplementation on the gut microbiota of C57BL/6 mice and this microbe’s effect on inflammatory responses after airway LPS challenge, with a focus on sex-dependent variations. For this experiment, we divided eight-week-old C57BL/6 mice into three groups, ensuring a balanced number of males and females in each cohort (**Fig 1A, Table S1**). The mice are described as belonging to three treatment groups: the Baseline group (green) was sacrificed without any antibiotic or LPS intervention, while the other two groups—PBS and *Bacteroides*—were subject to a three-week antibiotic course followed by gavage with either PBS (light blue) or *Bacteroides* (orange). The *Bacteroides* gavage was comprised of a pool of ∼1 x 10^9^ CFU/ml of bacteria composed of previously published strains *B. dorei* CFPLTA003_2B, *B. thetaiotaomicron* ATCC 29741 and *B. fragilis* AD126T 3B^1^. These strains were chosen for their ability to downregulate the pro-inflammatory cytokine IL-8 in IL-1β−stimulated Caco-2 intestinal epithelial cells ^1^. Two weeks after gavage, we administered LPS nasally and sacrificed the mice 24 hours later. To account for cage effects, we evenly distributed the treatment groups across different cages. Throughout the study, we regularly collected stool samples for 16S rRNA gene amplicon sequencing and culturing to assess bacterial community composition.

**Figure 1.**
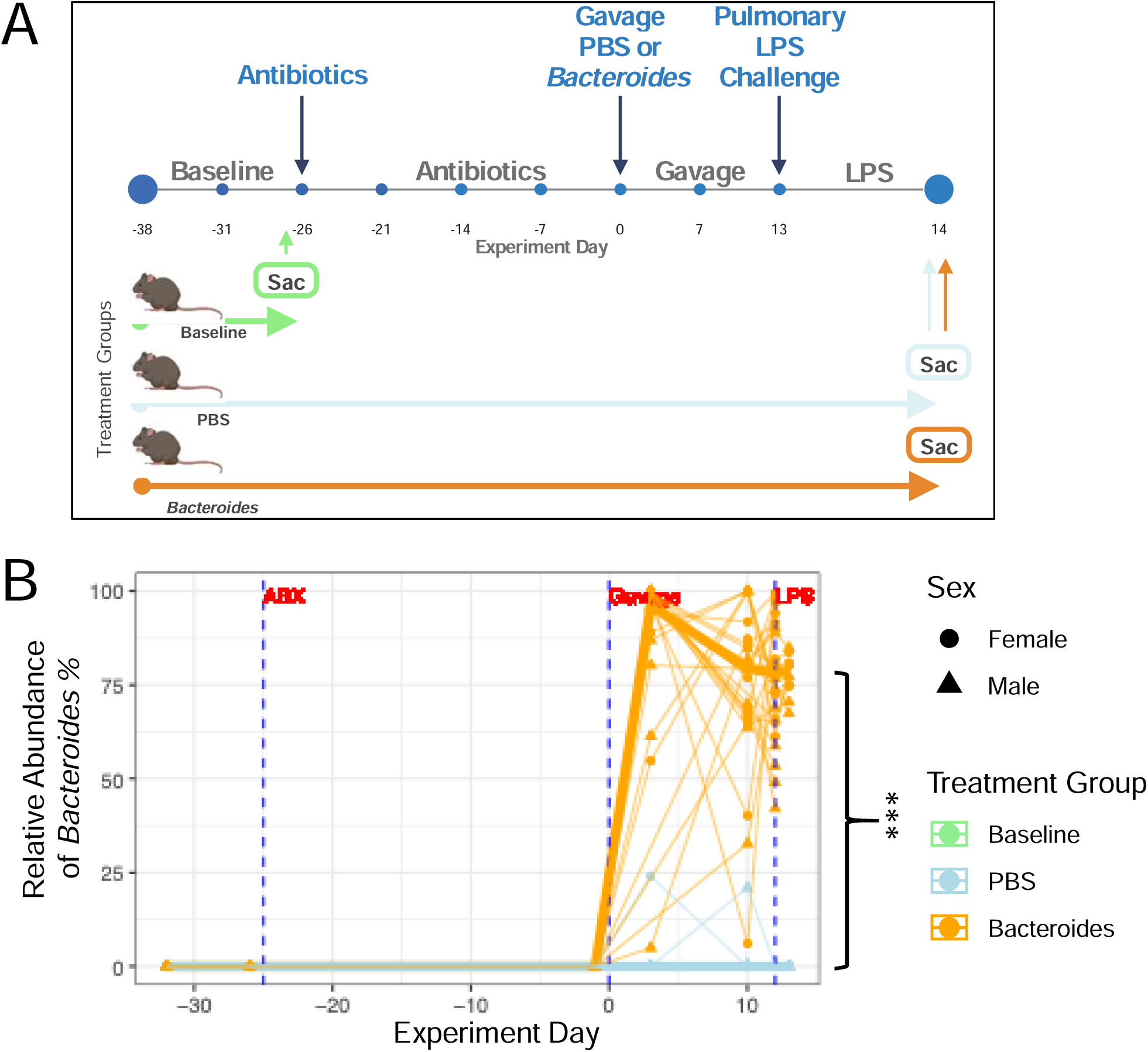
Experimental design and longitudinal analysis of gut microbiota following *Bacteroides* colonization and pulmonary LPS challenge. **A)** A schematic timeline detailing procedures performed on three groups of C57BL/6 mice, aged 8-12 weeks, balanced for sex. The baseline group (light green) did not receive any treatment and were sacrificed at 2 weeks post-baseline. The PBS (light blue) and *Bacteroides* (orange) groups were given antibiotics for three weeks to deplete their gut microbiota, followed by PBS or *Bacteroides* gavage. Two weeks post-gavage, an LPS challenge was administered nasally, and the mice were sacrificed 24 hours post-LPS challenge. This experiment was conducted twice (n=50). **B)** Stool samples were collected for 16S rRNA sequencing throughout the experiment to analyze gut microbiota changes over time. Symbols represent individual samples, with biological sex denoted by circles for females and triangles for males. The thinner lines show individual mouse sample progression, with bold lines illustrating average changes per treatment group. Statistical differences in *Bacteroides* relative abundance in post-LPS stool samples were assessed using a linear model with treatment group, sex, their interaction, and cage as fixed effects, enabling us to evaluate treatment and sex influences while accounting for cohousing effects. The *Bacteroides* treatment group had significantly greater relative abundance (p=2.97e-13), and the interaction between sex and treatment group showed that male mice had significantly decreased relative abundance of *Bacteroides* (p=0.0265, **Fig S1**) compared to female mice. ***, p < 0.001.

Our sequencing analysis revealed that *Bacteroides* has a significantly higher relative and absolute abundance in the stool of *Bacteroides*-gavaged mice versus PBS gavaged mice (**Fig 1B, Fig S1A**), and that sex of mouse influenced relative abundance of *Bacteroides* – this microbe establishes significantly less in males versus females (**Fig S1**). Sex of mouse did not influence absolute abundance of *Bacteroides* (**Fig S1B**). Stool samples were also plated on blood agar supplemented with 100 μg/mL gentamicin, a concentration to which *Bacteroides* exhibits inherent resistance, and grown anaerobically. This selective plating allowed us to monitor the presence and proliferation of *Bacteroides* in the gastrointestinal tract of the mice throughout the experiment (**Fig S2A**).

To assess microbial community richness and evenness, we measured alpha diversity of stool samples collected throughout the experiment (**Fig S3A**). Interestingly, gavage with *Bacteroides* is associated with an increase in Fisher alpha-diversity in stool starting soon after gavage, and this difference persists until LPS treatment, at which point there is no difference in alpha diversity between the treatment groups (**Fig S3A**). Analysis of Observed Richness (left) and Pielou’s Evenness (right) alpha diversities revealed that differences are driven more by richness than evenness (**Fig S3A**, inset).

Stool samples were also plated on MacConkey agar then incubated under aerobic conditions to track the viability of enteric organisms, including *Escherichia* spp., throughout the experiment, given the observation that this microbe is enriched in the CF gut^8,12^. Mice gavaged with *Bacteroides* showed significantly reduced CFU/mL on MacConkey medium (**Fig S4A**). However, there were no significant differences in relative abundance of *Escherichia-Shigella*, between the treatment groups across the experiment (**Fig S4B**), across biogeography (**Fig S4C**), or microniche (**Fig S4D**). Together, these data show that *Bacteroides* spp. introduced by gavage can allow this microbe to establish in the microbiota-depleted conventional mouse, impact the diversity of the intestinal microbial community, and potentially reduce the burden of facultative anaerobes.

### Systemic and airway inflammatory regulation by *Bacteroides* is sex dependent

Our previous studies showed that introduction of *Bacteroides* into the gut of an antibiotic-treated, conventional CF mouse resulted in lower airway KC levels upon pulmonary LPS challenge compared to a PBS control^1^. This work helped to establish the concept of the gut-lung axis in CF. To confirm systemic and distal inflammatory responses post-LPS challenge in this experiment after introduction of *Bacteroides* by gavage, and effectively validate our earlier studies in CF mice, serum and lung tissue samples were analyzed using a 32-plex Mouse Luminex panel. As expected, analysis comparing systemic and pulmonary cytokine levels between nonCF and CF mice at baseline showed a significant increase in KC and other proinflammatory cytokines in CF mice (**Fig S5A-B**).nonCF mice gavaged with *Bacteroides* exhibited significantly lower systemic KC concentrations (**Fig 2A**) compared to the PBS treatment group. Both sex and cage conditions notably influenced systemic KC levels. Mice gavaged with *Bacteroides* also demonstrated reduced lung KC level compared to the PBS group (**Fig 2B**). The combined effect of sex and treatment impacted pulmonary KC levels as well (**Fig 2B**). Thus, these studies are consistent with our previous work in CF mice, while noting a clear difference in response of the mice based on biological sex. Intriguingly, this protective effect of *Bacteroides* was more pronounced in female mice, which exhibited a more notable reduction in serum (reduction of ∼370 pg/mL in females vs ∼320 pg/mL in males) and lung (reduction of ∼420 pg/mL in females vs ∼245 pg/mL in males) KC levels compared to their male counterparts (**Fig S6A-C**).

**Figure 2.**
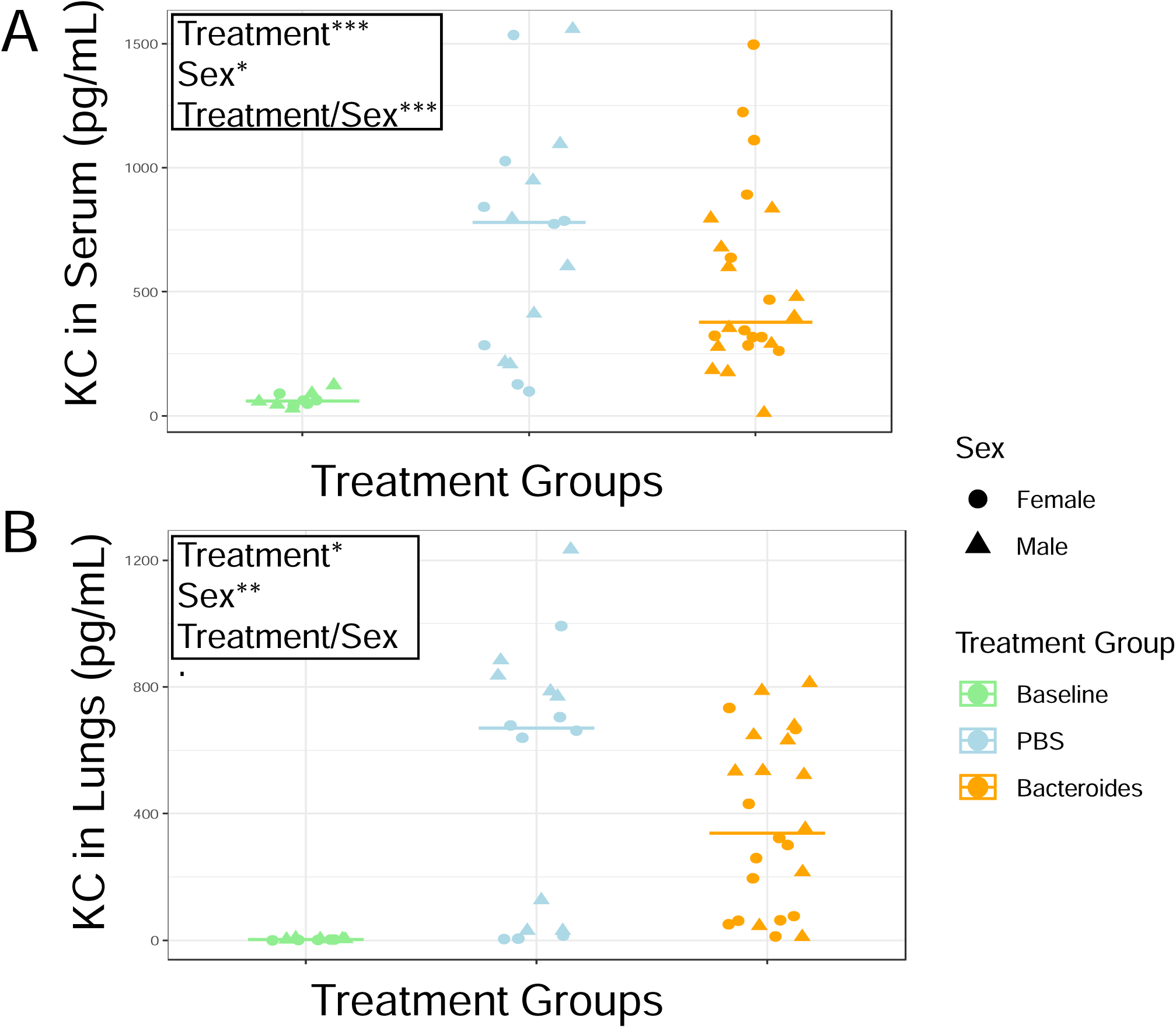
*Bacteroides* colonization attenuates systemic and pulmonary KC-mediated inflammatory responses in a sex-dependent manner following pulmonary LPS challenge. **A)** Serum and **B)** lung KC cytokine levels were measured, using a Luminex 32-panel assay, to assess the proinflammatory cytokine response post-LPS challenge. Statistical analysis was performed using linear models on logL-transformed KC values to evaluate the effects of treatment, sex, their interaction (Treatment/Sex), and cage (to account for cohousing). P-values for fixed effects are shown in the top left of each plot comparing PBS and *Bacteroides* Treatment groups. Significance: ‘ ‘, P > 0.05;., P < 0.1; *, p < 0.05; **, p < 0.01; ***, P < 0.001.

We next examined the correlation between serum and airway KC levels to better understand the relationship between systemic and pulmonary inflammation regulated by *Bacteroides* spp. in the gut. Interestingly, there is a significant positive correlation between serum and airway KC levels for the animals treated with antibiotics and gavaged with PBS. In contrast, the *Bacteroides*-gavaged animals showed a consistent level of systemic cytokine levels across a range of airway KC level (**Fig S7**). We discuss this point further below.

We further analyzed a broader panel of cytokines to identify those differentially regulated by *Bacteroides* colonization in both serum (**Fig S8A**) and lung tissue (**Fig S8B**). While KC was significantly downregulated in mice gavaged with *Bacteroides*, we observed that VEGF was significantly upregulated in these same animals in the serum. This contrasting regulation pattern suggests that *Bacteroides* selectively modulates specific inflammatory and growth factor pathways rather than causing a uniform suppression of immune signaling. Interestingly, we also see that mice gavaged with *Bacteroides* had significantly greater weight gain after gavage (**Fig S9**), indicating that this microbe might also impact the host’s ability to effectively acquire intestinal nutrients.

### *Bacteroides* establishes in the large intestine and predominates in the lumen

To understand where in the intestines *Bacteroides* is establishing in this mouse model, we collected luminal contents and the mucosal tissue of various regions across the intestine (**Fig 3A**). Intestinal segments and microniches (lumen and mucosa) were plated on blood agar + 100ug/mL gentamicin in anoxic conditions to measure viable bacteria. These same samples were extracted for DNA and subjected to 16s rRNA gene amplicon sequencing to determine relative abundance of bacteria. Amplicon sequencing revealed the highest relative abundance of *Bacteroides* in the lumen of the colon, reaching up to 65% in the proximal colon and 70% in the distal colon of the *Bacteroides-*gavaged mice (**Fig 3B**). The median relative abundance in the cecum is 38%. In contrast, the small intestine showed minimal colonization at a median of 5% (**Fig 3B**). These findings align with previous studies showing *Bacteroides* tends to flourish in the distal colon where conditions are more anoxic^6,13,14^. When comparing the relative abundance of *Bacteroides* across microniches, *Bacteroides* has a significantly higher relative abundance in the lumen than on the mucosal surface (**Fig 3C**).

**Figure 3.**
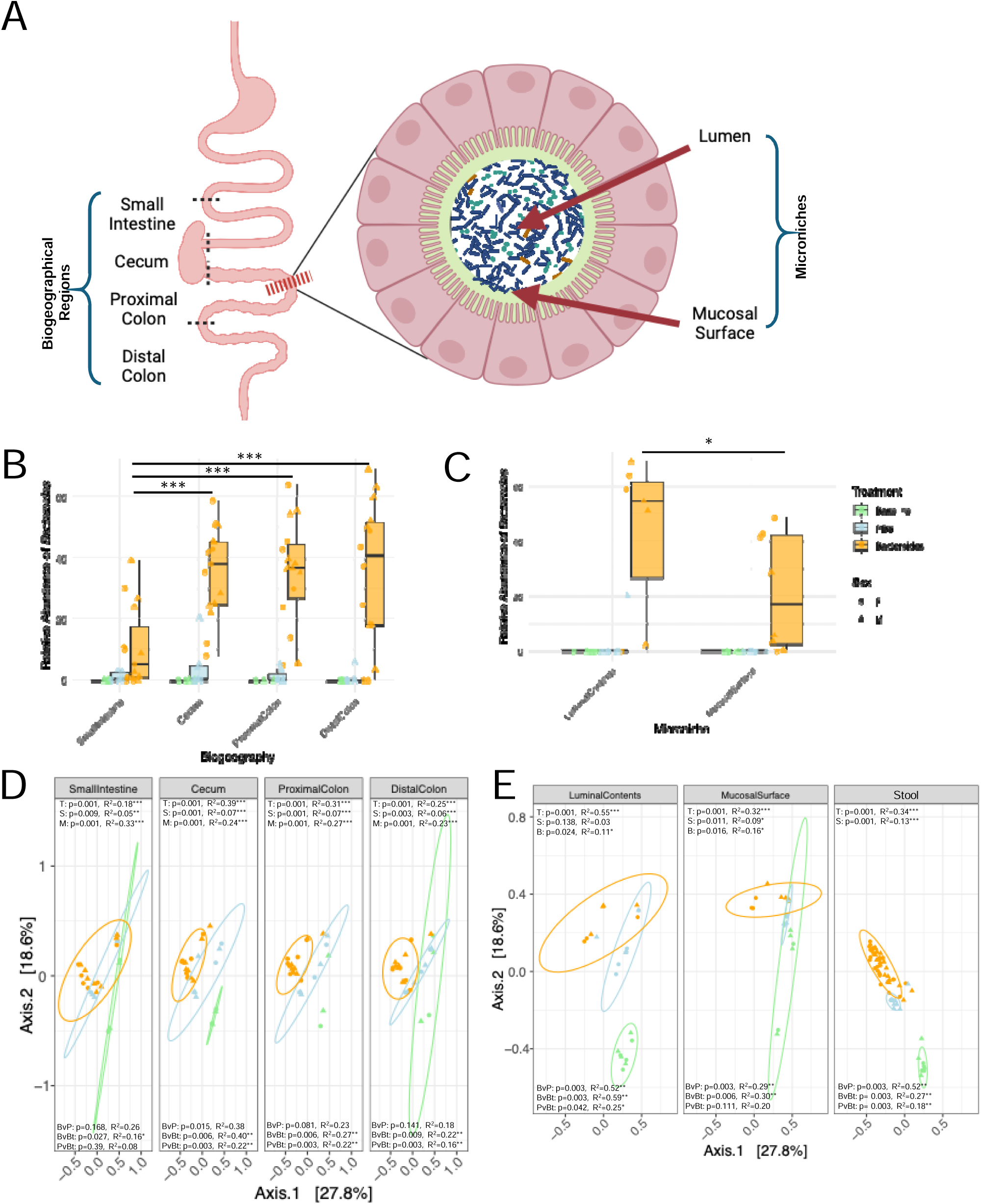
*Bacteroides* establishes in the large intestine and predominates in the lumen, driving community composition in these regions and niche. **A)** Shown are the anatomical divisions of the mouse intestine used for microbiota analysis. The intestinal regions isolated by incisions made along the dotted lines are the small intestine, cecum, proximal colon and distal colon. For each intestinal region, two microniches were examined (inset indicated by the red dashed line): the luminal contents and the mucosal surface. **B)** 16s rRNA gene amplicon sequencing is performed on DNA from each of the biogeographical and microniche regions to assess *Bacteroides* establishment across the gut. Treatment groups are color-coded: baseline (light green), PBS (light blue), and *Bacteroides* (bright orange), with biological sex represented by shapes: female (F; circle), male (M; triangle). A linear model was performed to determine the effect of biogeographical region on *Bacteroides* relative abundance. The small intestine was set as the reference, and significance was found within the *Bacteroides* treatment group, where the Cecum (p=0.0001), Proximal Colon (p=0.0001), and Distal Colon (p=0.0001) all had significantly more *Bacteroides* than the small intestine – however no difference between the three regions of the large intestine, as determined by linear model. **C)** Comparing *Bacteroides* relative abundance between the microniches, the mucosal surface shows lower relative abundance of *Bacteroides* than the lumen (p=0.0149). Beta diversity was assessed using Bray–Curtis distances to evaluate shifts in microbial composition across **D)** biogeographical regions and **E)** microniches. Statistical significance was determined by PERMANOVA, accounting for treatment (T), sex (S), their interaction, microniche (M) or biogeographical region (B), while accounting for cohousing and repeated measures. P-values for fixed effects appear in the top left of each PCoA plot. Pairwise comparisons between treatment groups (e.g., Baseline vs. PBS = BvP; Baseline vs. *Bacteroides* = BvBt; PBS vs. *Bacteroides* = PvBt) are shown in the bottom left. Significance: ***, P < 0.001; **, P < 0.01; *, P < 0.05; ‘ ‘ = not significant.

We next explored how *Bacteroides* might be influencing the ecology of microbial communities across the gut that rebounded after antibiotic treatment stopped. To address this question, we analyzed Fisher alpha diversity, comparing microbial communities of mice gavaged with PBS vs *Bacteroides* across the gut (**Fig S3B**) and microniches (**Fig S3C**). In this next analysis we include the Baseline group, to compare alpha diversity across the intestine prior to any kind of treatment with antibiotic or gavage. Our results revealed significant differences in baseline alpha diversity among the various gut regions. Specifically, the cecum and proximal colon displayed significantly higher alpha diversity (Fisher) compared to the small intestine at Baseline. Both PBS and *Bacteroides* treatments showed lower alpha diversity (Fisher) in the large intestine compared to Baseline. There was no difference in alpha diversity across all treatments in the small intestine (**Fig S3B**).

A similar analysis was conducted comparing the microniches of the intestines. There was no significant difference in alpha diversity (Fisher) between the mucosal surface and luminal contents at baseline, however both PBS and *Bacteroides* treatments had significantly lower alpha diversity (Fisher) in both microniches (**Fig S3C**). Analysis of Observed Richness and Pielou’s Evenness reveal that the differences across the biogeographical regions (**Fig S3B**, insets) are driven more by richness than evenness, while driven by both across the microniches (**Fig S3C**, insets).

Evaluation of beta diversity using Bray-Curtis distances revealed that treatment had a significant impact on the microbial community composition across all gut regions (**Fig 3D**), explaining the greatest variation in the cecum (R^2^=0.39) and proximal colon (R^2^=0.31). Interestingly, sex also influenced microbial composition in all regions, although this factor accounted for only ∼ 5-7% of the variation. Microniche contributed to significant community differences throughout the intestine, with the strongest effect observed in the small intestine (R^2^=0.33). To investigate the role of *Bacteroides* in shaping gut microbial composition, we conducted pairwise Adonis comparisons between PBS and *Bacteroides* treatment groups. These analyses revealed that *Bacteroides* induced the most pronounced shifts in community composition in the large intestine (**Fig 3D**), whereas no significant difference was observed in the small intestine between PBS and *Bacteroides* groups. This finding is in agreement with the enhanced colonization of *Bacteroides* in the large intestine (**Fig 3B**). However, in the small intestine, *Bacteroides* exhibited significant differences compared to community composition at baseline, indicating the combined treatment of antibiotics and *Bacteroides* gavage drove community restructuring in this region, as would be expected.

We conducted a similar analysis to examine the effects of Treatment (i.e., PBS and *Bacteroides* groups) on the beta diversity using Bray-Curtis distances for microniches and stool (**Fig 3E**). We found that Treatment drives significant changes across all three microniches, explaining more variation in the luminal contents (R^2^=0.55). Interestingly, sex plays a significant role in community composition in stool and at the mucosal surface with a low explanation of variation (R^2^=0.13 and 0.09, respectively), however not in the luminal contents. Overall, the pairwise adonis test shows that *Bacteroides* gavage drives significant differences in the luminal contents and stool after antibiotic treatment, but not at the mucosal surface (**Fig 3E**).

Our investigation across various biogeographical regions and microniches of the mouse intestine revealed that *Bacteroides* successfully establishes, predominantly in the luminal region of the large intestine. Consistent with this finding, the mucosal surface appears less affected by *Bacteroides* treatment, suggesting microniche-specific effects across the gut. Overall, these findings emphasize the spatially distinct influence of the introduced *Bacteroides* on gut microbiota across intestinal regions and microniches.

### Differential gene expression analysis for *Bacteroides*-versus PBS-gavaged mice across intestinal regions

To explore the impact of *Bacteroides* on host response throughout different regions of the intestine we conducted a comparative analysis between mice gavaged with *Bacteroides* vs. PBS. The central question we addressed was how the introduction of *Bacteroides* influences gene expression across the gut’s biogeographical regions. To this end, we extracted RNA from dissected regions of the intestine (**Fig 3A**, biogeographical) and subjected this RNA to NanoString Host Response gene set analysis, which provided a profile of gene expression patterns relevant to host immune response, antiviral response, host susceptibility and homeostasis across 785 genes (**Fig S10, Table S2**). We applied mixed-effects models to account for treatment, sex, and their interaction, allowing us to pinpoint genes/pathways in which expression was significantly altered by *Bacteroides*.

Our main findings highlight a regional variation in the response to *Bacteroides* (**Fig 4A** and **Table S2**). The small intestine exhibited the largest number of significantly differentially expressed genes (DEGs), with sixty genes showing significant differential regulation by *Bacteroides* compared to the PBS treatment group. This response was modestly less pronounced in the cecum (n = 55 DEGs) and further muted in the proximal and distal colon (n = 23 and 26 DEGs, respectively). These results suggest that *Bacteroides* elicits a strong localized response in the small intestine, which may not be as prominent in the colon.

**Figure 4.**
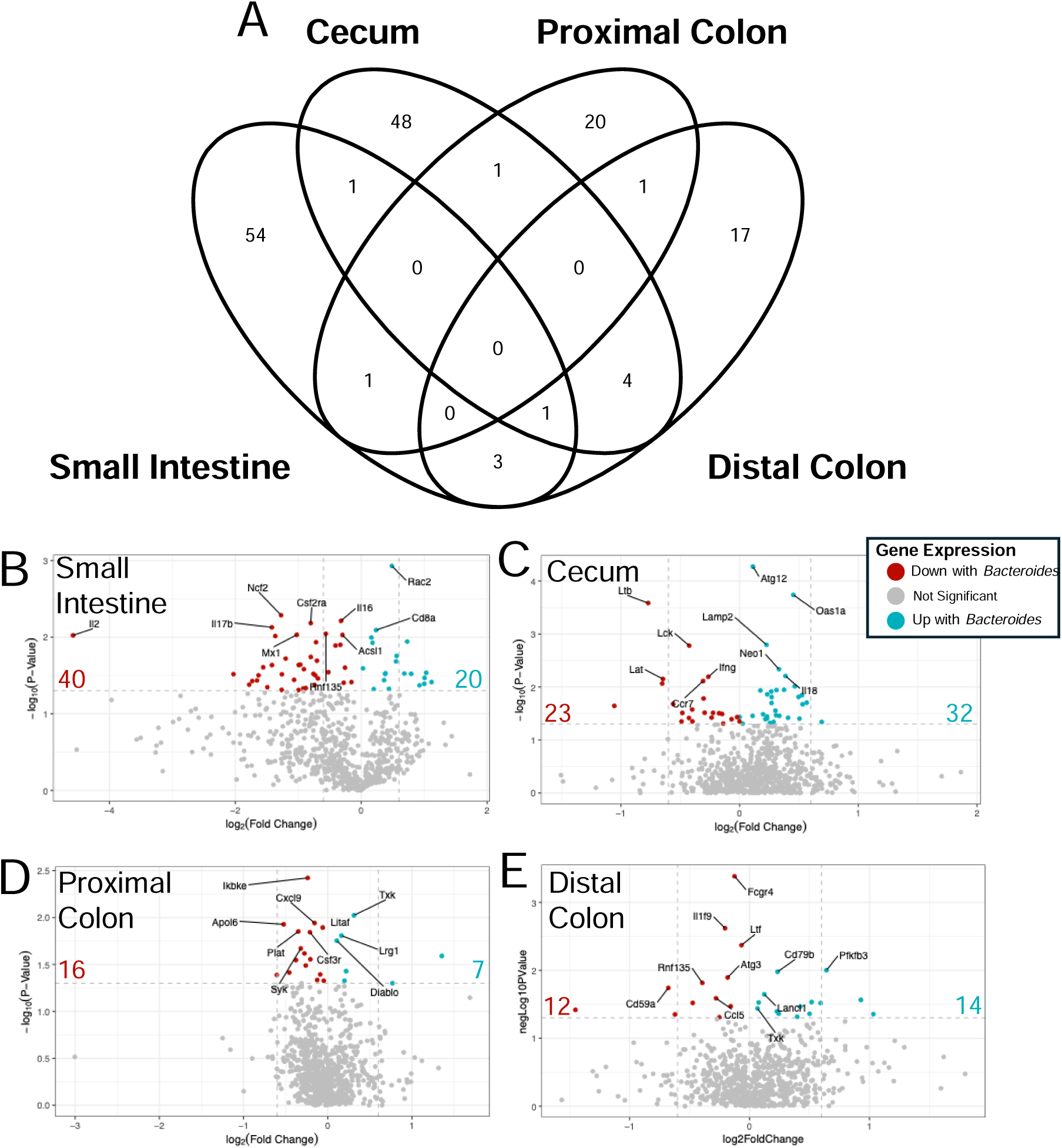
Differential gene expression in *Bacteroides* vs PBS treated mice across intestinal regions. **A)** A Venn diagram that illustrates the overlap and unique genes with significant differential expression (p-value ≤ 0.05) across different regions of the mouse intestine (small intestine, cecum, proximal colon, and distal colon) in response to *Bacteroides* versus PBS gavage. Each section of the diagram represents genes specific to, or shared among, these biogeographical regions. The colors dark blue, royal blue, light blue, and lightest blue represent the small intestine, cecum, proximal colon, and distal colon, respectively. Numbers inside the diagram indicate the count of significantly differentially expressed genes in each section. A volcano plot that presents the differential gene expression analysis between PBS and *Bacteroides* gavaged mice in the **(B)** small intestine, **(C)** cecum, **(D)** proximal colon, and **(E)** distal colon. Numbers on each side of the volcano plot indicate number of total genes up (in blue) or down (in red) significantly differentially expressed in the *Bacteroides* treatment group in that region, compared to the PBS treatment. Top ten most significantly regulated genes are labeled by gene name. Significant gene expression (p-value) was determined by performing a mixed-effects model on the logL transformed counts of each gene within each region, accounting for treatment, sex, and the combined effect of sex and treatment. Fold change was calculated by dividing the median of the PBS group by the median of the *Bacteroides* treatment group, per gene.

We further determined the number of genes that are up- vs. down-regulated in the presence of *Bacteroides* in each region, and found that *Bacteroides* was associated with more downregulation of genes in the small intestine (**Fig 4B**) and proximal colon (**Fig 4D**), while this microbe was associated with more up-regulation of genes in the cecum (**Fig 4C**) and distal colon (**Fig 4E**). When we performed pathway analysis of the DEGs (**Fig 5**), we noted that the presence of *Bacteroides* showed an enrichment for the adaptive immune response (n = 82), innate immune response (n = 73), interferon response (n = 38 genes), homeostasis (n = 33) and host susceptibility (n = 4) across all regions.

**Figure 5.**
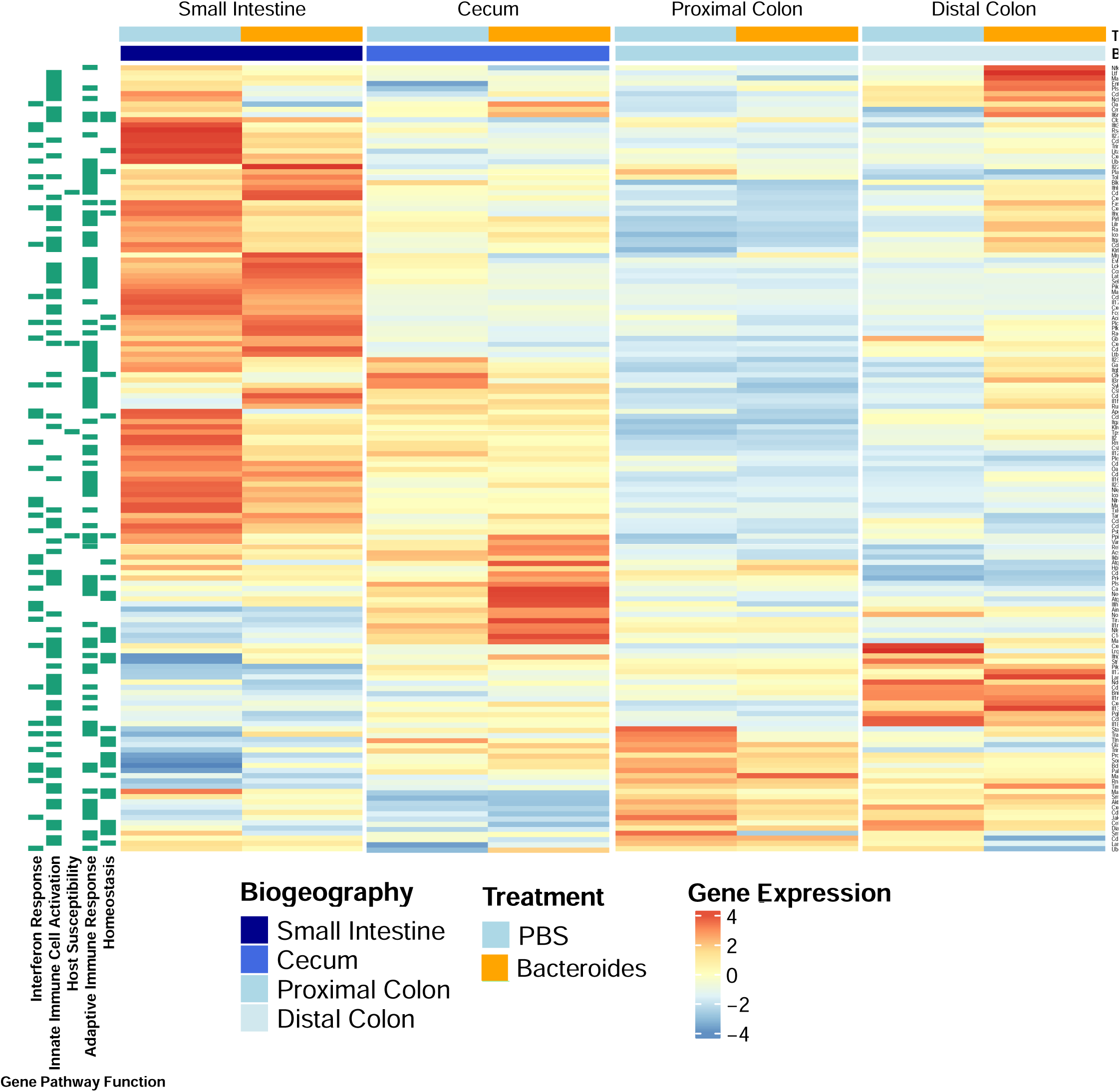
Heatmap depicting differential gene expression across mouse intestinal regions post-Bacteroides or PBS gavage. A heatmap illustrates the genes significantly differentially expressed across various biogeographical regions of the mouse intestine, comparing *Bacteroides* to PBS gavage. Each row of the heatmap corresponds to a specific gene, while columns represent the average gene count per biogeographical region (small intestine, cecum, proximal colon, and distal colon) and treatments (PBS and *Bacteroides*). Rows are scaled to show relative expression levels of each gene across different conditions. Genes in each row are further annotated according to their NanoString functional pathways (**Table S4**). The color intensity indicates the magnitude of gene expression: red for higher expression and blue for lower expression. Column annotations denote the biogeographical region and treatment type. The heatmap is ordered to cluster genes with similar expression patterns.

### Transcription factor (Txk) regulating G-protein coupled receptors is differentially expressed in the proximal and distal colon by *Bacteroides*

Because *Bacteroides* was predominately establishment in the large intestine, we first analyzed which genes were most significantly regulated across both the proximal and distal colon. When investigating genes differentially regulated by *Bacteroides* (compared to PBS) in both colonic regions, one gene was significantly upregulated in both locations: *txk* (**Fig 6A**). This gene encodes a nonreceptor tyrosine kinase transcription factor whose expression was previously shown to be negatively correlated with methylation patterns in human inflammatory bowel disease (IBD) blood samples^15^. *Txk* is also known to regulate G-protein coupled receptor 31 (GPR31) in humans, with a correlation value of 0.96 (The Human Protein Atlas)^16^. GPR31 has previously been shown to respond to short chain fatty acids and play a role in protecting host tissue from other pathogenic microbes, including *Salmonella*^17^. Statistical analysis of *txk* expression by mixed effects model revealed significant increase in expression in *Bacteroides*-treated animals in the proximal and distal colon (**Fig 6A**), where the combined effect of treatment and sex had significant effects in the distal colon.

**Figure 6.**
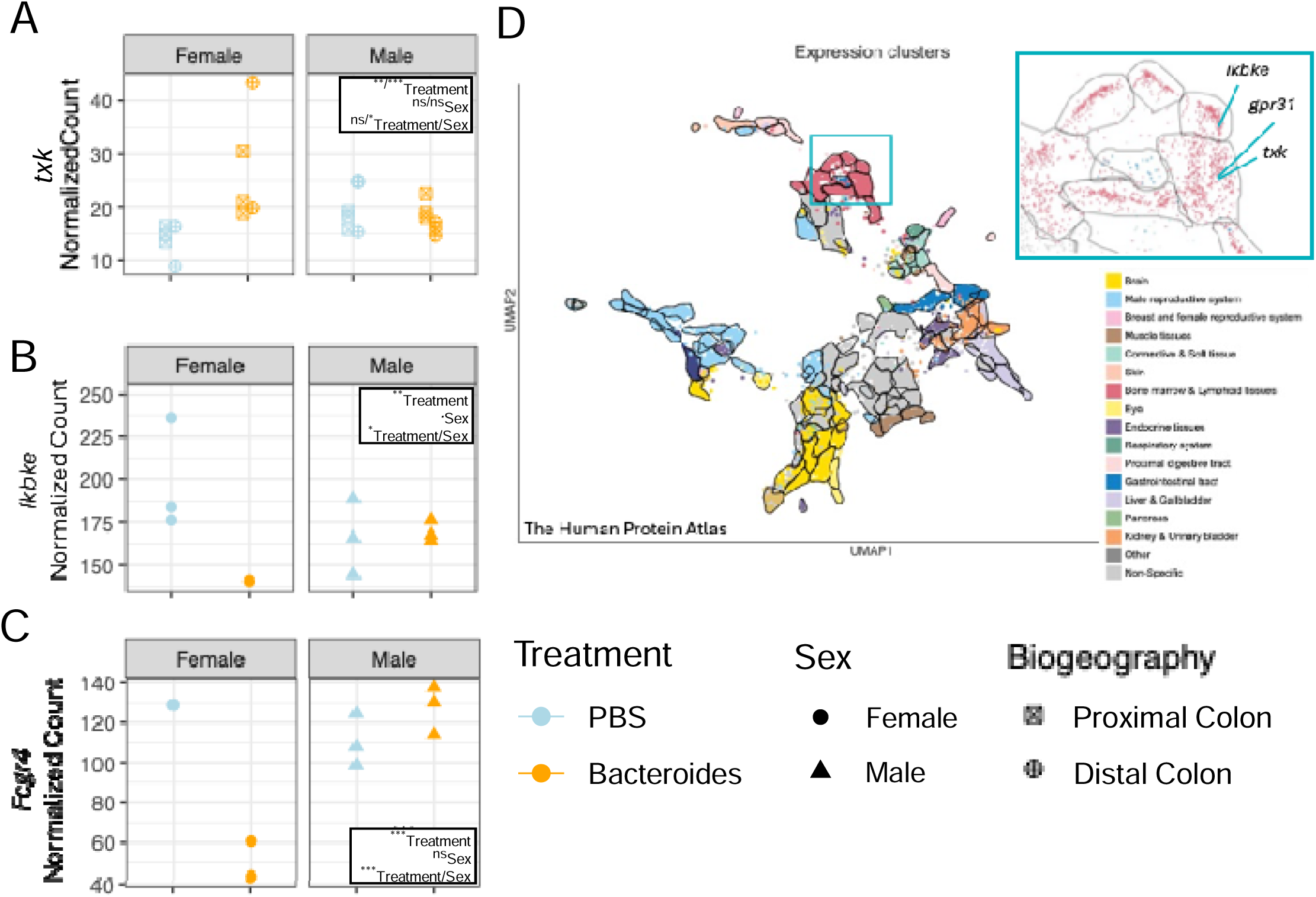
A Transcription factor (*txk*) for G-protein coupled receptors is differentially expressed in the proximal and distal colon post-*Bacteroides* gavage. **A)** The *txk* gene was the only gene significantly regulated by *Bacteroides* in both the proximal and distal colon— regions where *Bacteroides* is most established - while other genes showed region-specific regulation. This figure compares *txk* expression between PBS- (blue) and *Bacteroides*-gavaged (orange) mice. Each point represents an individual mouse, with biogeography of sample source indicated by shape (proximal colon, square and distal colon, circle). Statistical analysis was performed using a linear model on logL-transformed normalized counts, with PBS as the reference. The model included fixed effects for treatment, sex, and their interaction. Statistical significance is reported as results for proximal colon / distal colon. The most differentially expressed gene in the proximal colon and distal colon were (B) *ikbke* and (C) *fcgr4*, respectively. Plots depict normalized gene count on the y axis for each treatment group on the x axis, stratified by sex. Statistical analysis was done by a linear model, with PBS as the reference treatment. The model was run on log2 transformed normalized counts in the proximal colon and accounts for the effects of treatment, sex, and their combined effect. Significance codes: ns, P > 0.05;., P < 0.1; *, p < 0.05; **, p < 0.01; ***, P < 0.001. **D)** The Human Protein Atlas revealed genes that are regulated by the transcription factor *txk* in humans, where RNA data was analyzed to group genes based on their similar expression patterns across various samples. Each cluster of genes, identified through this analysis, was categorized and colored based on shared functions and specific traits. The top right panel shows a zoom in on those genes differentially expressed by *Bacteroides*.

The genes with most significantly differential expression in the proximal and distal colon, respectively, are *ikbke* (inhibitor of nuclear factor kappa-B kinase subunit epsilon), an oncoprotein to whose high expression is associated with cancer cell growth and is a target for cancer therapeutics in colon and lung cancers^18,19^ (**Fig 6B**), and *fcgr4*, which is an analog of the MS4A2 protein in humans and promotes IgE-induced lung inflammation^20–22^ (**Fig 6C**), respectively. Our findings revealed significant effects in the proximal colon by treatment (p=0.0095), however sex had no significant effect (p=0.0770). The model for the distal colon revealed treatment (p=0.0041) and the combined effect of sex and treatment to be significant (p=0.0007). Importantly, analysis of the Human Protein Atlas revealed that the transcription factor *txk* in humans impacts the expression of *ikbke* and *fcgr4* (**Fig 6D**), consistent with the activation of this pathway in the intestine upon introduction of *Bacteroides.* These results suggest that this microbe has the capacity to modulate immune pathways throughout the intestine while also exerting distinct influences in specific regions of the gut.

### Expression dynamics of *atg12* and macrophage-related genes in response to *Bacteroides* gavage in the mouse cecum

In our analysis, *atg12* emerged as the most significantly upregulated gene in the cecum following administration of *Bacteroides*, compared to the PBS control group (**Fig 7A**). Notably, female mice displayed a pronounced increase in *atg12* expression, whereas males experienced a modest decrease. This gene is central to the autophagic process^23^, a critical cellular mechanism for maintaining equilibrium, recycling cellular debris, and orchestrating immune functions, including the function of macrophages^23–25^, a point we further explore in the Discussion. Given the connection of this gene to macrophage function^26,27^, we further investigated the genes included in the panel that are markers for macrophages: *cd163* (**Fig 7B**), *cd68* (**Fig 7C**), *cd84* (**Fig 7D**), and *ms4a4a* (**Fig 7E**). *Bacteroides* administration significantly increased the expression of macrophage markers *cd163*, *cd84*, and *ms4a4a*, with a trend toward increased *cd68* expression, suggesting an alteration in the macrophage biology in the cecum compared to the other regions of the gut.

**Figure 7.**
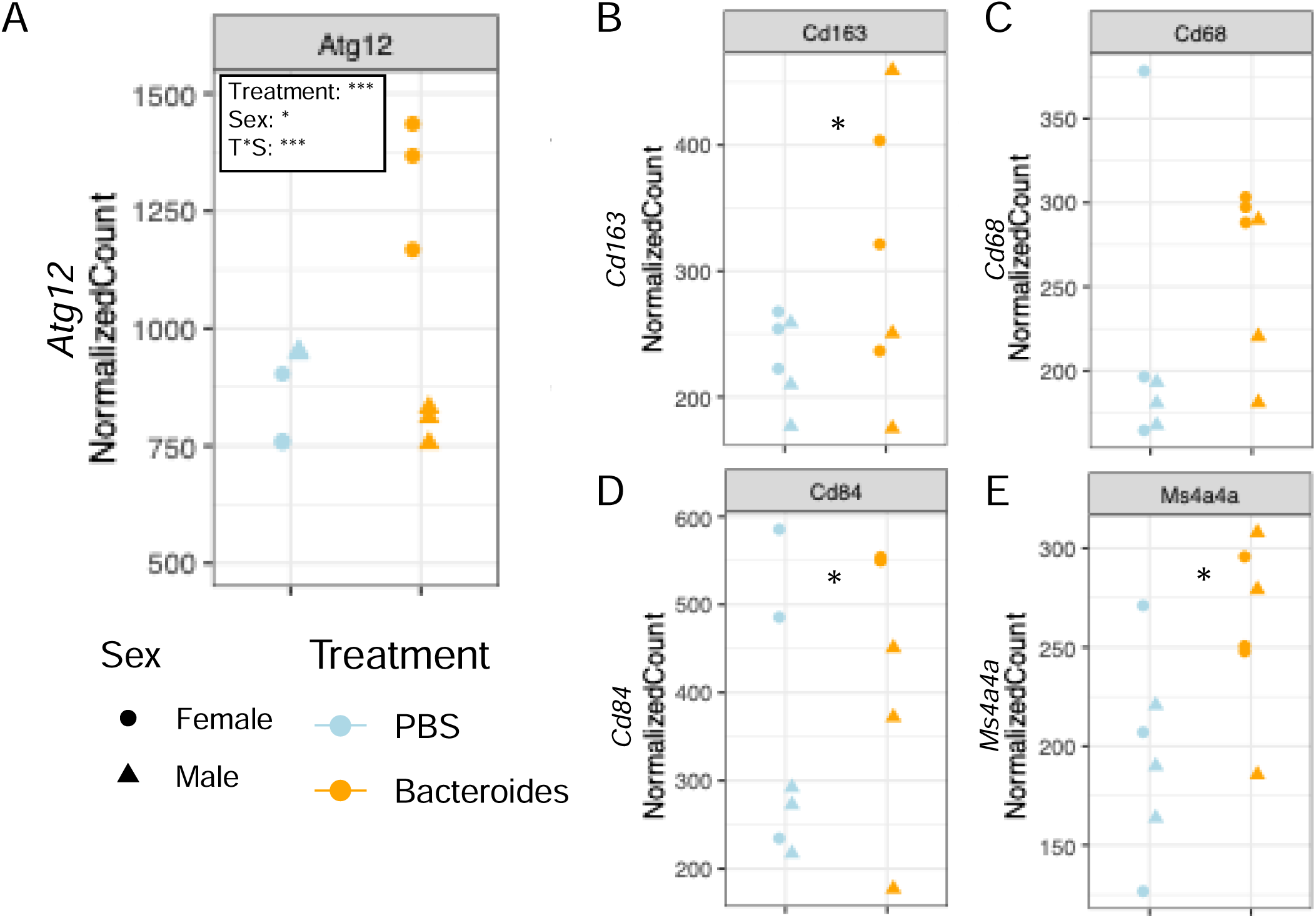
Expression dynamics of *atg12* and macrophage-related genes in response to *Bacteroides* gavage in mouse cecum. **A)** The plot illustrates *atg12* expression in the cecum of mice gavaged with PBS (blue) or *Bacteroides* (orange). Individual data points reflect each mouse, with males as triangles and females as circles. The graph captures the variation in *atg12* expression post-gavage and LPS challenge. Statistical analysis was performed with a linear model, setting PBS as the reference treatment. The models were run on log2 transformed normalized counts and accounts for the effects of treatment, sex, and their combined effect. P-values for fixed effects are shown in the top left of each plot. Subsequent panels focus on the expression of macrophage-associated genes **B)** *cd163*, **C)** *cd68*, **D)** *cd84*, and **E)** *ms4a4a.* Mixed-effects linear models were used to analyze log_2_-transformed normalized counts for each gene, accounting for treatment, sex, biogeography, and repeated measures from mice. Significance codes: ***, P < 0.001; **, P < 0.01; *, P < 0.05; ns = not significant.

## Discussion

Our study investigates the systemic and gut-specific responses of mice to intestinal *Bacteroides*, incorporating the variable of biological sex. We aim for these findings to shed light on the broader ecological implications of intestinal microbial supplementation by *Bacteroides*, as well as the intricate immune responses that follow in each region, and how these immune responses may play a role in the gut-lung axis. Our findings verify the successful colonization of the intestines by *Bacteroides* post-antibiotic treatment and gavage. Additionally, we confirm that *Bacteroides* administration via oral gavage offers a protective effect against systemic and lung-specific proinflammatory responses triggered by a pulmonary LPS challenge in WT mice, as evidenced by the levels of the cytokine KC measured in serum and lungs.

To explore how gut-resident *Bacteroides* may influence systemic versus localized inflammatory responses, we examined the correlation between serum and airway KC levels following LPS challenge. In antibiotic-treated, PBS-gavaged mice subsequently challenged with LPS in the lung, there was a strong positive correlation between serum and airway KC levels (R^2^=0.87), indicating a coordinated systemic and pulmonary inflammatory response. However, in mice colonized with *Bacteroides*, serum KC levels remained relatively stable and lower regardless of variation in airway KC levels (R^2^=0.03). These findings suggest that *Bacteroides* colonization in the gut exerts a stabilizing effect on systemic inflammation, potentially creating a buffering mechanism that prevents excessive inflammatory responses from propagating throughout the body. While LPS challenge still induces detectable increases in serum and pulmonary KC levels, *Bacteroides* appears to establish a regulatory threshold that limits the magnitude of this systemic response. This compartmentalization of inflammation may explain the protective effects observed in distal organs, as perhaps *Bacteroides* prevents localized inflammatory triggers from cascading into a widespread inflammatory state.

Analysis of stool samples through 16S rRNA sequencing revealed that in some mice in our studies, *Bacteroides* constituted up to 100% of the gut microbiota. We further wanted to understand where in the intestines *Bacteroides* is establishing and to assess how the host responds in each region. We observed the highest relative abundance of *Bacteroides* in the large intestine, comprising 38-41% in the cecum, proximal colon, and distal colon. In contrast, the small intestine showed minimal colonization, at approximately 5% relative abundance. The counts of *Bacteroides* as assessed by gentamicin resistant colonies is ∼10-fold lower in the small intestine as compared to the large intestine, indicating that there is a relatively low level of the microbe persisting in this region of the gut in our studies. *Bacteroides* are more abundant in the luminal contents across the intestine, comprising 55% of the microbial populations, while their relative abundance drops to 17% at the mucosal surface. Notably, mice gavaged with *Bacteroides* exhibited a significant reduction in CFUs/mL on MacConkey agar under oxic conditions, suggesting that the introduction of *Bacteroides* to the gut may modulate nutrient availability and perhaps occupy a niche that could otherwise be dominated by other Gram-negative bacteria.

Sex has also been shown to play a role in how diet impacts gut microbiota. One study saw that a high fat diet increases abundance of *Alistipes*, *Lachnospiriceae*, *Lactobacillus*, and *Clostridium* in male mice but not female mice^28^. In their analysis of how different patient characteristics influence the beta diversity of gut microbiota in a cohort of children with CF, Price et al. discovered that sex significantly impacts the beta diversity in children with CF aged 0-2 years, along with age, breastfeeding, and delivery mode^29^. However, these characteristics only explained ∼8% of the variability across all samples. Here we see that sex has a significant impact on the relative abundance of *Bacteroides* after gavage, with females exhibiting 15% higher median relative abundance of *Bacteroides* after LPS challenge compared to male mice. Notably, no difference was observed in the absolute abundance of *Bacteroides*, indicating that sex influences the overall community composition rather than the colonization of *Bacteroides* in the gut. Further research is essential to comprehend the influence of sex on gut microbiota.

The pattern of differentially expressed genes (DEGs) across the intestine highlights the most substantial changes occurring in the small intestine (n = 60 DEGs), followed by a moderately lower level in the cecum (n = 55) and more pronounced reduction in the number of DEGs in the proximal (n = 23) and distal colon (n = 26). The observation that largest number of differentially expressed genes was in the small intestine, where *Bacteroides* relative abundance were lowest is a surprising result. It is possible that we missed a narrow but critical window for capturing *Bacteroides* populations in the small intestine as we assessed gene expression ∼2 weeks after the *Bacteroides* gavage. Alternatively, even a brief transit of *Bacteroides* through the small intestine might be sufficient to elicit the observed host response. Lastly, it is conceivable that impacts of *Bacteroides* in the proximal and distal colon were sufficient to elicit a broad host response across the intestine, the pronounced changes observed in the small intestine may reflect its higher density of immune cells compared to other intestinal regions^3^.

We noted that the only gene for which there was a significant regulation dependent on the presence of *Bacteroides* in both the proximal and distal colon, was *txk*, which encodes a transcription factor reported to have high correlative transcription with G coupled receptor 31 (GPR31) in humans (The Human Protein Atlas^16^). GPR31 is known to respond to short chain fatty acids, such as those shown to be secreted by the *Bacteroides* strains used here^61^ and that were shown to be essential for protective host response. GPR31 plays an essential role oral tolerance or averting mucosal tissue-related disorders like food allergies, celiac disease, and IBD, as well as systemic immune tolerance induced by microbiota secretions within the gut ^30^.

In the proximal colon, a notable decrease was observed in the expression of *ikbke* in female mice gavaged with *Bacteroides*, while no change was detected in males. *Ikbke* encodes IKBKE (inhibitor of nuclear factor kappa-B kinase subunit epsilon), a key regulator of immune signaling and cell proliferation. IKBKE has been implicated as an oncoprotein, with elevated expression linked to the progression of colon and lung cancers and proposed as a potential therapeutic target^18,19^. In addition to its role in tumorigenesis, IKBKE functions in antiviral defense and immune regulation via its interaction with TBK1, facilitating the production of interferons essential for antiviral immunity^31^.

In the distal colon, our analysis showed a significant decrease in the expression of *fcgr4* in female mice gavaged with *Bacteroides*. The *fcgr4* gene is identified by NanoString as a marker for neutrophils; this gene encodes the fragment crystallizable receptor of IgG, low affinity IV FCGR4 in mice. The Fcgr4 receptor resembles the human MS4A2 protein, which promotes IgE-induced lung inflammation. The FCGR4 receptor is involved in IgG2b-mediated inflammatory responses^32^, and when deleted, there is a reduction in neutrophil recruitment^33^. The reduction of this gene’s expression associated with *Bacteroides* may implicate a reduction in neutrophil recruitment to the distal colon whose regulation is specific to female mice.

In the cecum, we identified *atg12* as the gene most significantly modulated by *Bacteroides*, with an increase in expression observed in females (by ∼40%) and a slight decrease in males (by ∼5%). *Atg12* plays a pivotal role in autophagy, a vital cellular process for discarding damaged organelles and modulating immune responses. This gene, as part of the autophagic machinery, partners with *atg5* to enable the formation and degradation of autophagosomes. Autophagy is particularly crucial in the gut for regulating epithelial cell turnover, mucosal immunity, and responses to microbial incursions.

In CF, both human and mouse macrophages display a diminished ability to eliminate bacteria like *B. cenocepacia*, *Pseudomonas*, *Staphylococcus*, and non-tuberculous *mycobacteria* from the airway. This reduction in bacterial clearance correlates with lowered autophagic activity in CF immune and epithelial cells. Autophagy-related genes such as *atg5*, *atg12*, and *atg7* are significantly under-expressed in CF macrophages compared to non-CF macrophages^27^. Our findings suggest that particularly in female mice, *Bacteroides* induces an upregulation of *atg12*, potentially enhancing autophagy function. This observation prompted further investigation into whether *Bacteroides* gavage influenced macrophage-related gene expression, leading to the discovery that three out of four macrophage markers found on the Nanostring panel (*cd163*, *cd84*, and *ms4a4a*) were differentially regulated in the *Bacteroides*-treated group within the cecum. These data suggest that the presence of *Bacteroides* in the gut could reverse the reduced ATG12 pathway function noted for CF, at least for females, and suggests a mechanism whereby *Bacteroides* in the gut could perhaps impact airway macrophage function, a hypothesis worthy of future study. Additionally, CD163 is a well-established marker of M2 macrophages, which are associated with anti-inflammatory and antimicrobial functions. In mouse models, deletion of *Cd163* leads to an exaggerated inflammatory response to allergic stimuli, yet these mice exhibit increased susceptibility to infection, highlighting CD163’s role in balancing immune regulation and host defense^34^. CD84 is a signaling molecule implicated in immune suppression; its upregulation has been linked to immunosuppressive environments, suggesting a potential role in dampening immune activity and promoting tolerance^35^. Similarly, MS4A4A has been shown to promote M2 macrophage polarization and impair CD8L T cell function. It is highly expressed in the tumor microenvironment of colon cancers, where it likely contributes to immune evasion and tumor progression by fostering an immunosuppressive milieu^36^. Analysis comparing systemic and pulmonary cytokine levels between nonCF and CF mice at baseline revealed a significant increase in KC and other proinflammatory cytokines in CF mice. Given the previously observed protective effects of *Bacteroides* gavage on systemic and pulmonary KC levels, as well as its regulation of macrophage markers, *Bacteroides* gavage in CF mice may enhance macrophage function.

Our study represents an exploration of how the host responds systemically and in specific gut regions to *Bacteroides*, with a particular focus on sex differences in mice. We aimed to illuminate the ecological impact of intestinal microbial supplementation with *Bacteroides*, as well as the complex immune responses that arise in different intestinal regions, potentially influencing the gut-lung axis. This comprehensive analysis across the intestine is novel in its approach and offers insightful data into the intricate dynamics of microbial manipulation in gut ecology and the corresponding immune responses. Key observations include the successful colonization of the intestines by *Bacteroides* post-antibiotic treatment, with a notable protective effect against inflammatory responses triggered by pulmonary LPS challenges. This effect varied between sexes, with female mice displaying a more significant reduction in systemic and pulmonary inflammatory markers. The study also highlights the substantial changes in gene regulation across the intestine induced by *Bacteroides*, notably in the small intestine, cecum, and colon. These changes point towards the complex interplay between transient microbial presence and host immune responses. Additionally, the study observes sex-specific effects in gene expression related to immune regulation and gut health, further underscoring the intricate dynamics of microbial-host interactions. Moving forward, studies of beneficial microbes should carefully assess their impacts at both the mucosal interface and luminal space, while taking into account sex, to fully capture their ecological and immunological roles. These findings are pivotal in advancing our understanding of gut microbiology and its impact on health and disease, opening new pathways for therapeutic interventions targeting gut microbiota.

## Acknowledgements and Funding

We would like to express our deep gratitude to the Cystic Fibrosis Foundation for their support through grant #OTOOLE22G0 and the NIH via R01-ES033988. Our work has also been significantly advanced by the generous funding from the PhD Innovation Program through NSF grant #2125733. The samples, facilities and expertise offered by Dartmouth’s Translational Research Core have been invaluable to our research (NIH/P30-DK117469). Additional support was provided by the CF Training Grant to KEB (T32HL134598) and the CFF RDP (BOMBER24G0). Special thanks are due to Zack Peters and Arianna Reuven for their collection of stool samples, and Alex Fu who helped extract DNA. We are also grateful for the assistance provided by the animal facility staff, especially Eric DuFour and Kathy Bennet, whose expertise in gavage and euthanasia procedures ensured the completion of our mouse experiments. Thank you to Andrew Calkins for conducting the Luminex panel and to Heidi Trask at GMBSR, who processed our RNA samples with NanoString. Figures 1A and 3A in this publication were created with BioRender.com.

## Materials and Methods

### Mouse studies

C57BL/6 mice, acquired from Case Western Reserve, were maintained on a ScottPharma LabDiet 5V75 with free access to food and under a 12-hour light/dark cycle. Mice were housed within same treatment condition, but in multiple cages per treatment group and sex to account for cage a confounding actor. The following mouse experiment was conducted twice, with 50 total mice across the two experiments. A 21-day antibiotic regimen, described in prior studies^1^, was employed to diminish native intestinal flora, involving 1 mg/mL antibiotic solutions in sterilized drinking water. The regimen included alternating antibiotic cocktails over three 7-day periods, interspersed with breaks featuring only sterile water for 2 days (except for the last break, in which only 1 day of sterile water is provided prior to gavage). The first and third periods utilized a mix of ampicillin, cefoperazone sodium, and clindamycin hydrochloride, while the second period comprised ertapenem, neomycin sulfate, and vancomycin hydrochloride. Following the antibiotic course, mice received oral gavages based on weight (10uL/g mouse weight). Treatments of either PBS or a *Bacteroides* cocktail, composed of previously published on strains CFPLTA003_2B, SMC7758, and AD126T 3B^1^, were administered via oral gavage. Strains were anaerobically cultured on blood agar plates for 48 hours, collected with sterile wooden stick, washed and resuspended in PBS for administration. The *Bacteroides* cocktail amounted to roughly ∼10^9^ CFU/mL, while the PBS control had no bacterial contamination, confirmed via culture.

Fecal samples were collected throughout the experiment for microbial composition analysis, using various agars for culturing and 16S rRNA gene amplicon sequencing (SeqCoast). The fecal pellets were also plated to determine total bacterial counts on blood agar + 100ug/mL gentamicin in anoxic conditions to assess counts linked to *Bacteroides* spp., and MacConkey agar in oxic conditions to monitor *E. coli* populations at 37°C.

Mice then underwent a non-invasive intranasal 0.2mg/kg of LPS (*P. aeruginosa*, Sigma) challenge and were euthanized 24 hours post-challenge. Mice were sacrificed with Euthasol 24 hours post-challenge and whole lungs and segmented intestinal tissues from the small intestine/ileum, cecum, proximal colon, and distal colon were harvested for further processing. Proximal and distal colons were isolated by measuring the full length of the large intestine, cutting it in half, and identifying the half closer to the cecum as proximal and the half closer to the rectum as distal. Samples for CFU analysis were stored on ice until plating the same day. Samples for NanoString analysis were immediately submerged into RNALater and placed on ice, then stored at -80’C until RNA could be extracted. Whole lungs and blood from the heart were collected at the time of sacrifice for systemic cytokine analysis by Mouse 32-Multiplex, where lungs were immediately suspended individually after dissection into 5 mL of tissue lysis buffer (11 mg collagenase IV, 2.5 μL DNase I, 250 μL fetal bovine serum (FBS), and 4.75 mL PBS). Lungs were suspended in the tissue lysis buffer, homogenized using an electric homogenizer for 30 seconds and then incubated at 37°C for 45 minutes. Twenty microliters of 0.5M EDTA was added, and the sample was vortexed. Lung samples were then centrifuged at 1,500 rpm for 5 minutes in a microfuge. One milliliter of supernatant from each sample was collected into a 1.5 mL Eppendorf tube, and then the Eppendorf tube was centrifuged at 15,000 rpm for 10 minutes at 4°C to pellet debris. Finally, 200 μL of supernatant was aliquoted into a 96-well round bottom plate and stored at −80°C until analysis for cytokine levels. An aliquot of the leftover tissue lysis buffer was also stored to serve as a blank control. Serum was separated by allowing samples to stand at room temperature for 1–2 hours and centrifuged at 1,500 × g 4°C for 10 minutes. The upper fraction of serum was removed and stored at −80°C prior to submission for Luminex. Luminex processing was also done for whole lungs and blood collected from CF mice (Cftr^em1Cwr^ (an F508del with a milder CF phenotype) at baseline.

### 16S rRNA gene amplicon sequencing

Fecal pellets and intestinal samples were processed using the Zymo Quick-DNA Fecal/Soil Microbe Miniprep Kit and an aliquot of DNA was sent in for 16s RNA amplicon sequencing with SeqCoast. For 16S V3/V4 amplicon sequencing, samples were processed utilizing the Zymo Quick-16S Plus NGS Library Prep Kit, with the addition of unique dual indexes for differentiation. The sequencing occurred on an Illumina NextSeq2000 system, employing a 600 cycle flow cell kit, yielding 2x300bp paired-end reads. To enhance base calling accuracy for low diversity libraries within patterned flow cells, a 30-40% PhiX control, which lacks indexing, was incorporated into the library mixture. Sequences were further merged and taxonomy assigned using the R package DADA2. Alpha diversity, beta diversity, and relative abundance quantification was done using the R package Phyloseq and plotted using the R package ggplot2.

### Absolute abundance quantification

Stools samples collected 13-14 days after mice were gavaged with PBS or *Bacteroides* were selected for absolute abundance quantification as previously described^1^. Briefly, twenty uL of ZymoBIOMICS Spike-In Control I was added to 10 mg of each stool sample prior to DNA extraction by the Zymo Quick-DNA Fecal/Soil Microbe Miniprep Kit and sent for 16s rRNA gene amplicon sequencing. Quantification of absolute abundance was completed as described in the ZymoBIOMICS Spike-in Control 1 protocol.

### Luminex Analysis

In this analysis, we examine counts from the Mouse 32-Plex Assay (Millipore Luminex MCYTMAG-70K-PX32), including: Eotaxin/CCL11, G-CSF, GM-CSF, IFN-γ, IL-1α, IL-1β, IL-2, IL-3, IL-4, IL-5, IL-6, IL-7, IL-9, IL-10, IL-12 (p40), IL-12 (p70), IL-13, IL-15, IL-17, IP-10, KC-like, LIF, LIX, MCP-1, M-CSF, MIG, MIP-1α, MIP-1β, MIP-2, RANTES, TNF-α, and VEGF. In R, we implement mixed effects linear models to evaluate the influence of treatment (PBS vs *Bacteroides*) or genotype (Baseline nonCF mice vs Baseline CF mice), sex, and cohousing on individual log2transformed cytokine expression levels. We perform Bonferonni multiple comparisons correction. For volcano plots, we find fold change by taking the median cytokine count in the mice gavaged with PBS vs *Bacteroides*, or in CF mice vs nonCF mice.

### NanoString analysis

RNA was extracted from the dissected intestinal regions using the RNeasy kit from Qiagen. Whole tissue samples were placed into pre-weighed 2mL Eppendorf tubes containing 1mL of RNA Extraction Buffer and homogenized for 30 seconds. A quantity of 30mg of the homogenate was then processed following the manufacturer’s instructions. Post-extraction, the samples were treated with DNAse (Turbo, Invitrogen) to remove any genomic DNA contamination and processed using the NanoString Host Response Panel. For initial normalization, NanoString’s nSolver platform was employed. This involved subtracting the values of negative controls from each sample, with a minimum count threshold established at 20. Any samples exhibiting over-saturation, as indicated by the positive controls, were temporarily excluded from the analysis to be diluted and re-assessed. The saturation threshold was set between 0.3 to 3. Samples were further normalized to the panel’s housekeeping genes, allowing for a control variance between 51-71%. Housekeeping genes deviating from this variance range were excluded from the normalization process. After normalization, the gene count data of the qualifying samples were imported into R for in-depth statistical analysis.

### Statistical analysis

A summary of all the statistical analysis performed in this report can be found in **Table S3**.

