## Supplemental Figures for "Sex and regional effects of *Bacteroides* in the gut"

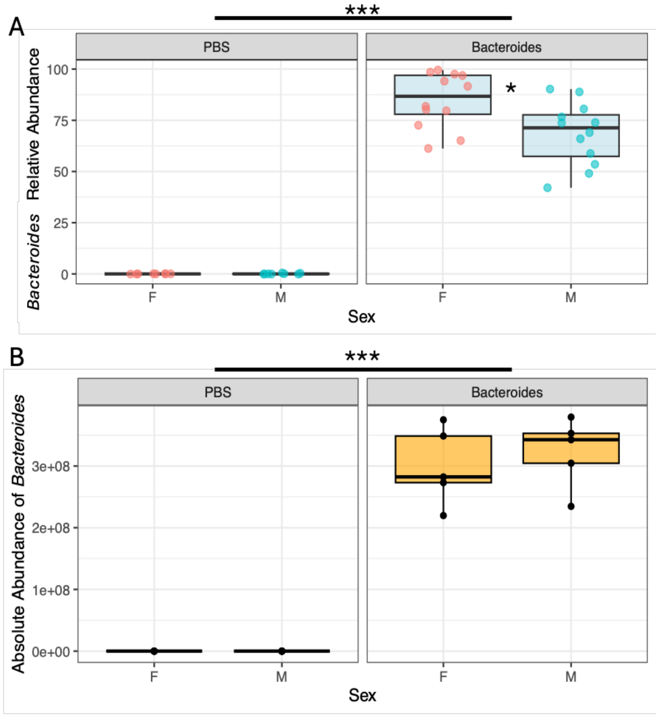


**Figure S1 *Bacteroides* establishes in greater relative abundance in female mice. A)** Stool samples were collected for 16S rRNA sequencing after LPS challenge, and relative abundance of *Bacteroides spp.* was determined and faceted by treatment group. Symbols represent individual samples, with sex denoted by red for females (F) and blue for males (M). Statistical differences in *Bacteroides* relative abundance in post-LPS stool samples were assessed using a linear model to evaluate treatment and sex influences while accounting for cohousing effects. While treatment showed significantly increased relative abundance of *Bacteroides* in the *Bacteroides* treatment group (p= 2.97e-13), the interaction between sex and treatment group showed that male mice had significantly decreased relative abundance of *Bacteroides (right panel* P=0.0265). **B)** Absolute abundance was measured by 16s rRNA gene amplicon sequencing from DNA extracted from the stool samples collected after LPS challenge, where a ZymoBIOMICSSpike-In Control I was added into each sample. Statistical differences in *Bacteroides* absolute abundance in post-LPS stool samples were assessed using a linear model to evaluate treatment and sex influences while accounting for cohousing effects. While treatment showed significantly increased absolute abundance of *Bacteroides* in the *Bacteroides* treatment group (p= 1.31e-05), the interaction between sex and treatment group showed that male mice had no difference in absolute abundance of *Bacteroides* (right panel P=0.713). ***, P < 0.001; *, P < 0.05; ns = not significant.

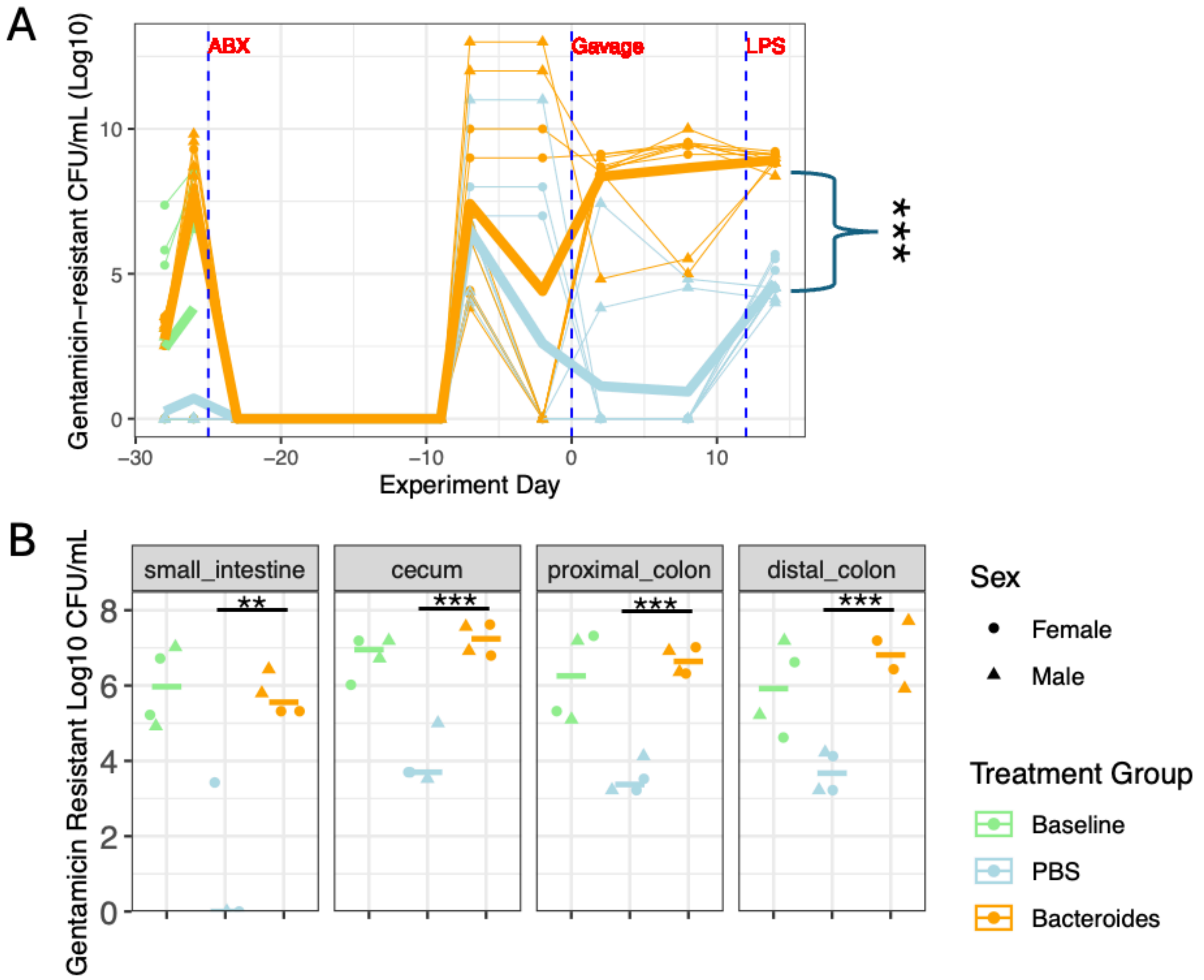


**Figure S2. Analysis of gentamicin-resistant CFU/mL in mice across experiment and biogeographical region. A)** Stool was collected throughout the experiment from mice, then serially diluted and plated to count CFU/mL on blood agar +100ug/mL gentamicin under anoxic conditions. Each point indicates stool collected from a single mouse, where a circle indicates female mice and triangle male mice. The thin lines connect stool collected from the same mouse, while the thick lines represent the mean of each treatment group across the experiment. Statistical significance was determined by mixed effects model on $log_{10}$ transformed CFUs/mL collected during the time-period labeled “gavage”, taking into account treatment, sex, their combined effect, and cage, as fixed effects and mouse ID and experiment day as a random effects to account for repeated measures from the same mice **B)** Luminal contents were collected at sacrifice from each treatment group and processed similar to stool. Statistical significance was determined by linear model for each biogeographical region, taking into account treatment and sex. Significance codes: ***, P < 0.001; **, P < 0.01; *, P < 0.05; ns = not significant.

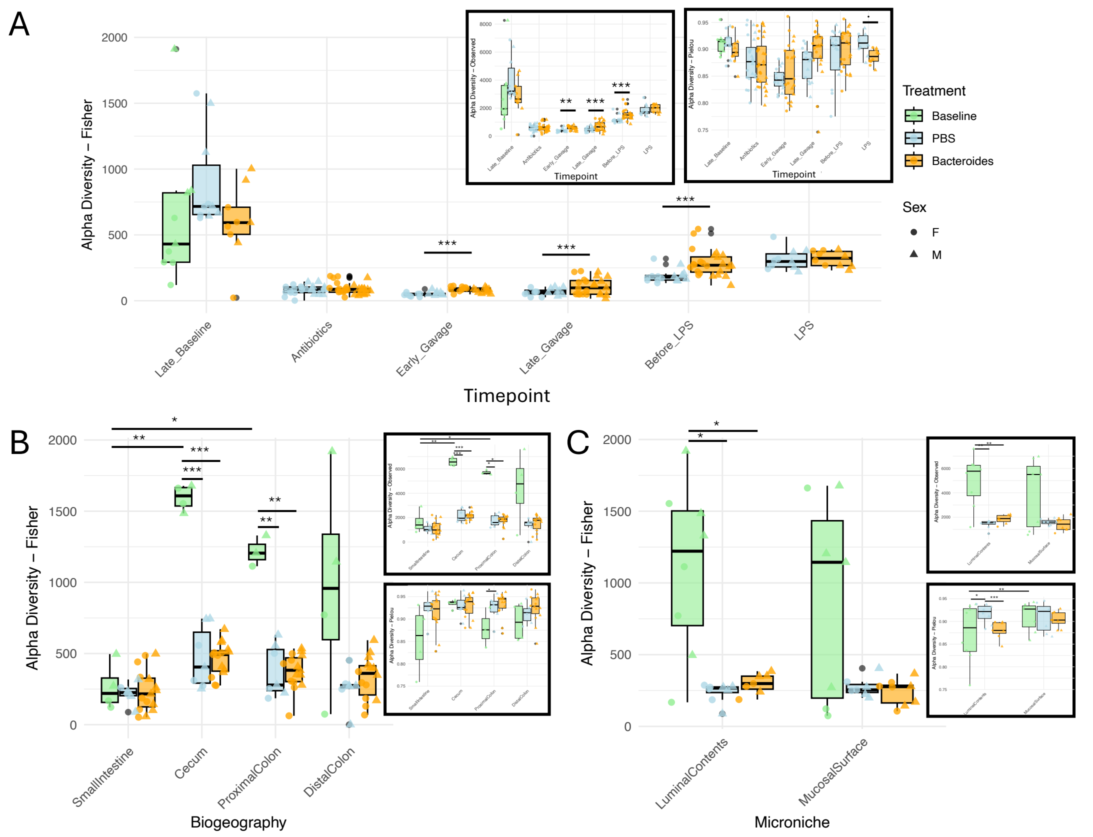


**Figure S3. Alpha diversity in stool across experiment and across biogeographical regions and microniches. A)** Fisher's alpha diversity of stool samples based on gut microbiota changes over time, colored by Treatment (Baseline, PBS and *Bacteroides*). Symbols represent individual samples, with gender denoted by circles for females (F) and triangles for males (M). Insets show Observed and Pielou's evenness at each time point. Significant differences between treatment groups were determined by linear model considering treatment, sex, their combined effects, and taking into account cohousing. Alpha diversity values were $\log_2$ transformed prior to running linear model if found to be nonnormal by Shapiro test. A similar analysis was done for each **B)** biogeographical region and microniche **(B)**. Further analysis was performed to determine how alpha diversity varies across the gut in different biogeographies and microniches at baseline. The results of a linear model reveal significantly higher Fisher's alpha diversity in the cecum (p=0015) and proximal colon (p=0.0429) as compared to the small intestine at baseline, taking into account biogeographical region, sex, their combined effects, and microniche region. In a similar analysis, Pileou's evenness was significantly greater at the mucosal surface (p=0.0014, inset) than the luminal contents at baseline. Significance codes: ***, P < 0.001; **, P < 0.01; *, P < 0.05; ns = not significant.

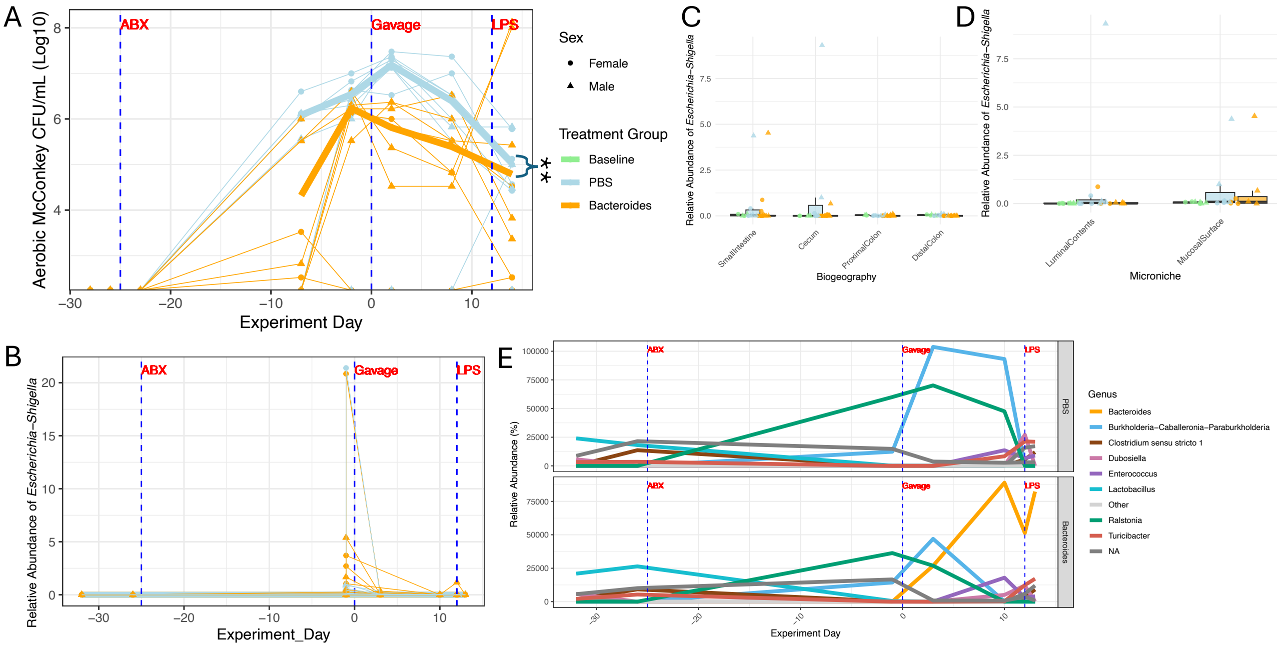


**Figure S4. *Bacteroides* gavage increases competitive inhibition of facultative anaerobes, but not specifically *E. coli*. A)** Stool was collected throughout the experiment from mice, then serially diluted and plated to count CFU/mL on McConkey under normoxic conditions. Each point indicates stool collected from a single mouse, where circle indicates female mice and triangle male mice. The thin lines connect stool collected from the same mice, while the thick lines represent the mean of each treatment group, indicated by color, across the experiment. **B)** DNA was extracted from stool collected throughout the experiment and analyzed by 16S rRNA gene amplicon sequencing. A) Relative abundance of *Escherichia-Shigella* in stool across the experiment**, B)** across the biogeographical regions and **D)** microniches of the gut. **E)** The relative abundance of the top 10 genera in stool samples collected across the experiment are plotted for PBS and *Bacteroides* treatment groups. Statistical significance was determined by mixed effects linear model on CFUs/mL collected after gavage, taking into account treatment, sex, their combined effects, and cage and accounting for time and cohousing. Significance codes: **, P < 0.01; ns = not significant.

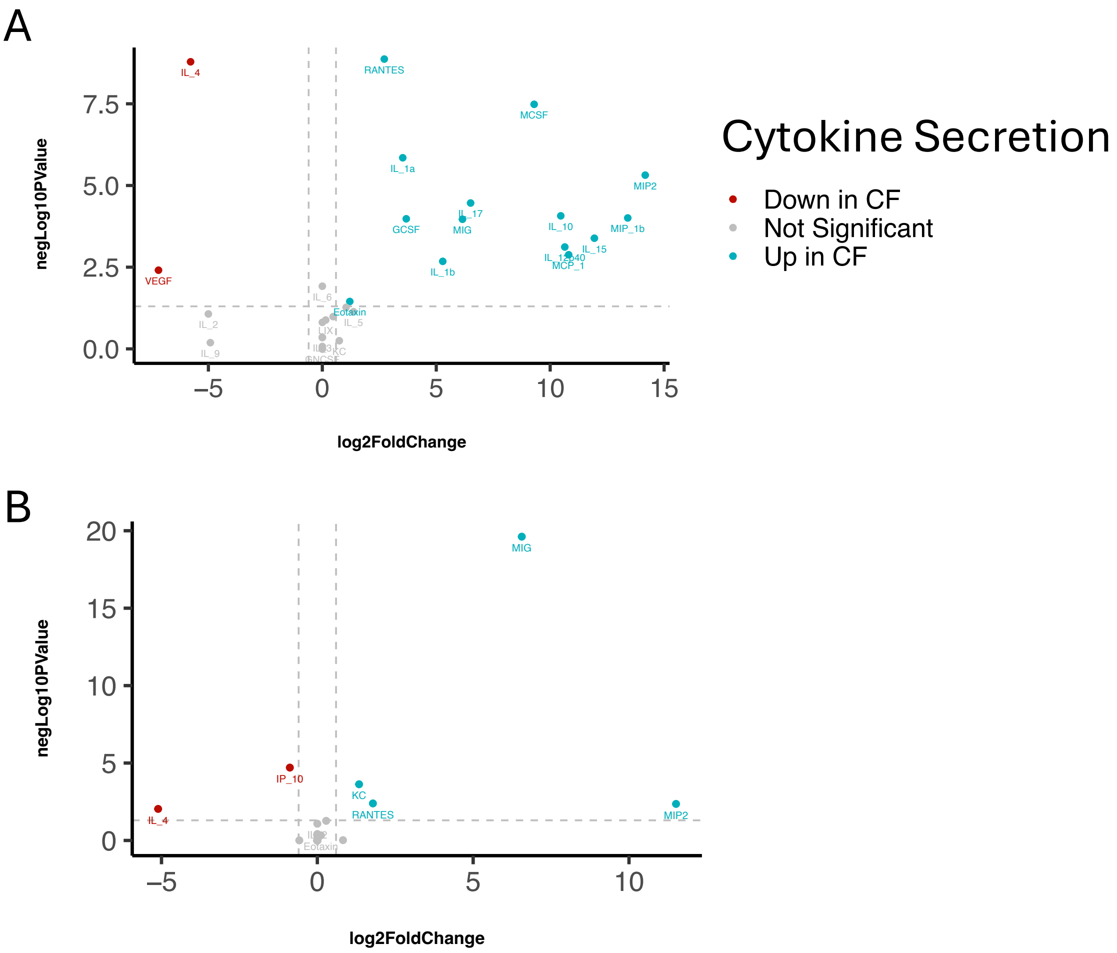


**Figure S5. Analysis of cytokine levels at baseline in nonCF and CF mice for A) serum and B) lungs.** For each cytokine from each source, the median value for nonCF and CF samples is determined, followed by computing the fold change as the ratio of the nonCF median divided by the CF median. Additionally, the $\log_2$-transformation of the fold changes is calculated. A linear model is constructed for each cytokine using genotype (nonCF or CF) and sex (male or female) to determine if they have a considerable effect on each cytokine level while considering cohousing. The negative $\log_{10}$ of the P-value from this analysis is plotted against the $\log_2$ fold change for the corresponding cytokine. In these plots, vertical dashed lines indicate a fold change cut off of 0.6, and horizontal dashed lines denote the p value cut off of 0.05. Notable findings include cytokines that are significantly down-regulated in CF mice (red) or up-regulated in CF mice (blue), while nonsignificant results are demonstrated in gray.

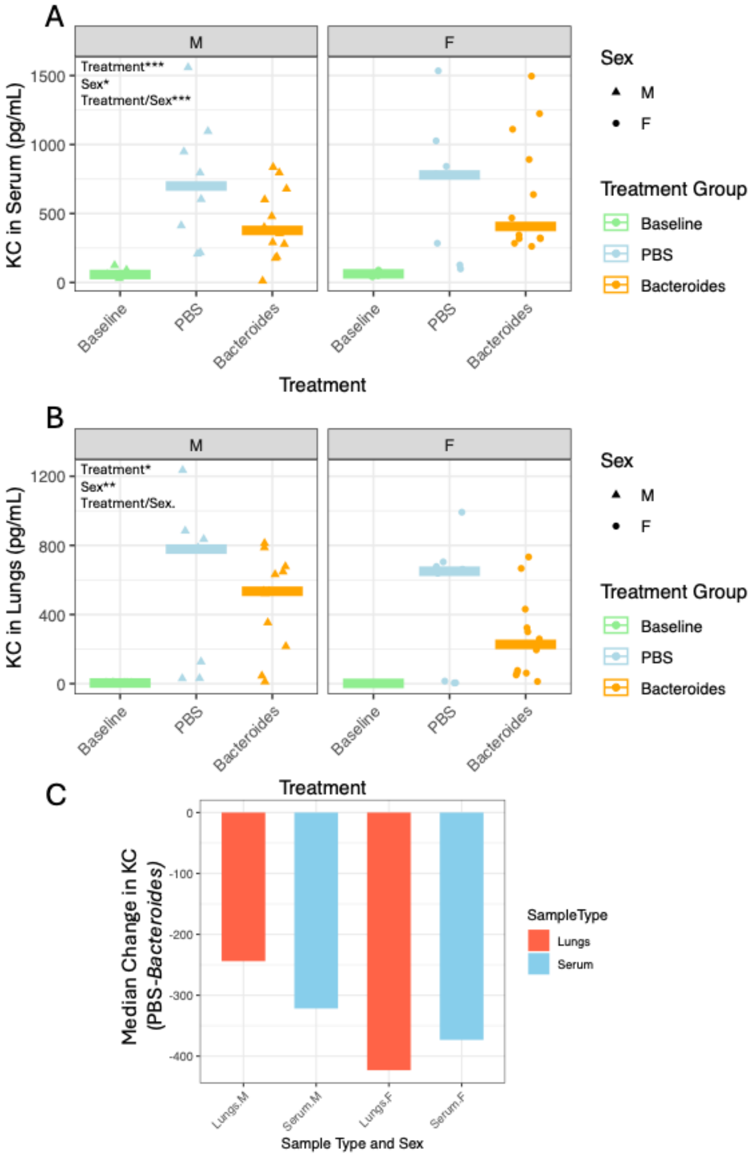


**Figure S6. *Bacteroides* colonization attenuates systemic and pulmonary KC-mediated inflammatory responses in a sex-dependent manner following pulmonary LPS challenge**
**A**) Comparison of KC levels in male and female mouse serum. **B**) Comparison of KC levels in male and female mice lungs. C) Visualizing the median difference between $\log_2$-transformed KC values for PBS vs *Bacteroides*-gavaged mice for each sex on the X-axis, with female mice displaying a more notable protective effect against increased systemic and pulmonary KC levels following LPS challenge. Statistical analysis was performed using linear models on $\log_2$-transformed KC values to evaluate the effects of treatment, sex, their interaction (Treatment/Sex), and cage (to account for cohousing). P-values for fixed effects are shown in the top left of each plot comparing PBS and *Bacteroides* Treatment groups. Significance codes: ***, $P < 0.001$; **, $P < 0.01$; *, $P < 0.05$; ns = not significant.

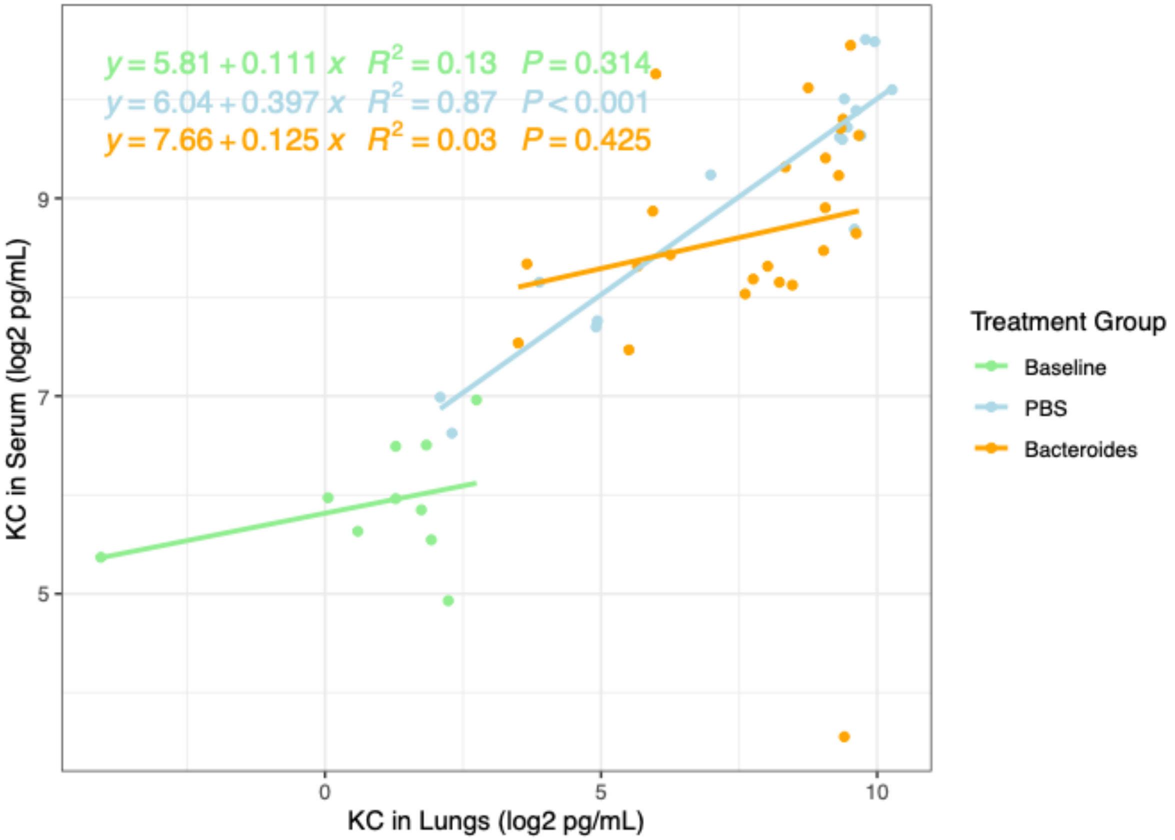


**Figure S7. Correlation of serum and lung proinflammatory KC secretion in mice post-*Bacteroides* gavage.** This figure presents a correlation analysis of the inflammatory marker KC levels in the serum and lungs of mice subjected to different treatments. Each data point corresponds to log-transformed KC levels from an individual mouse, with serum levels plotted on one axis against lung levels on the other. A line of best fit is drawn for each treatment group, illustrating the relationship between systemic and localized inflammation. The line fit, $R^2$ value and P value are inset in the figure panel and color-coded to the corresponding line color.

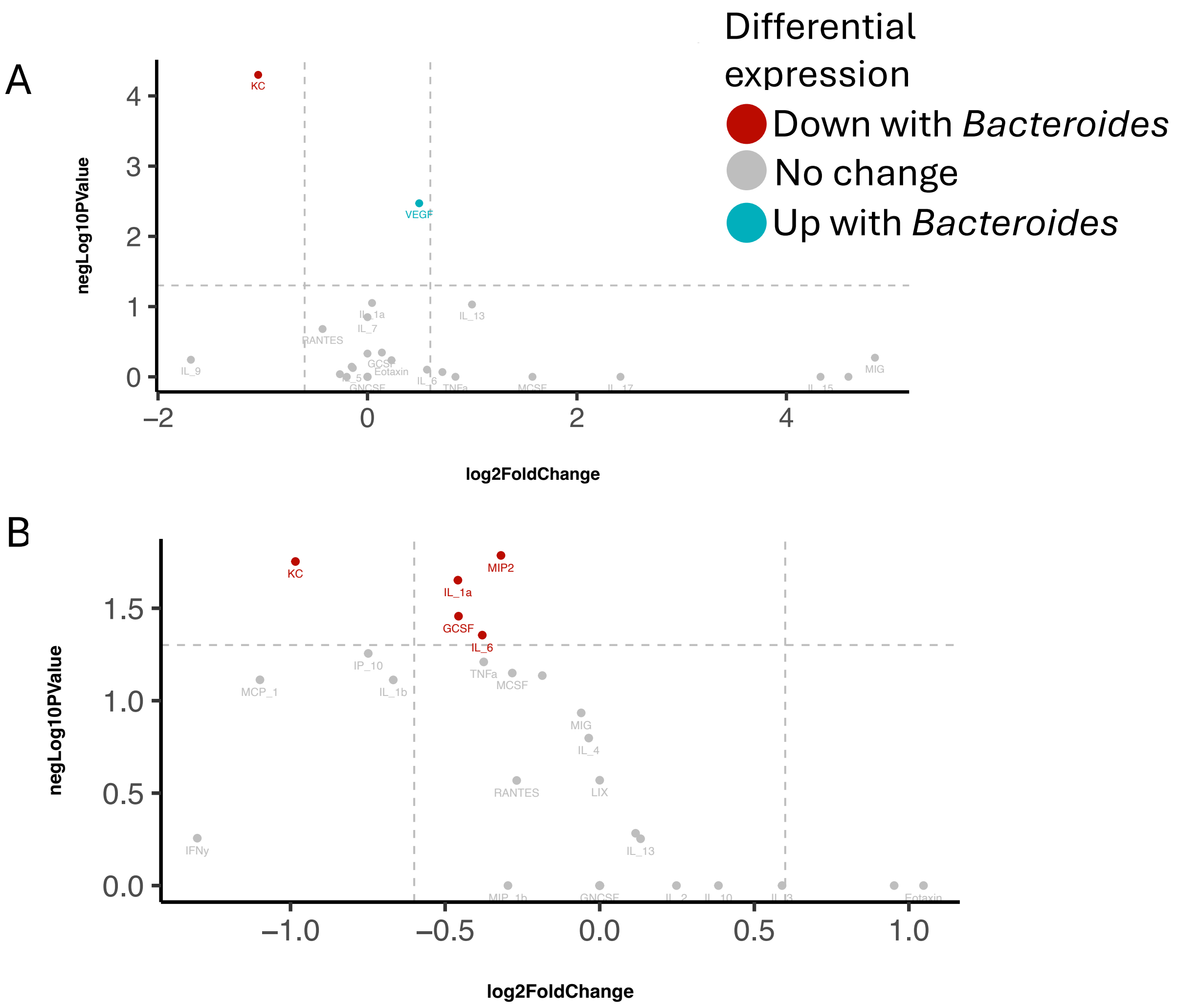


**Figure S8. Analysis of a cytokine panel in the serum and lung.** We further analyzed the broader Luminex panel of cytokines to identify those differentially regulated by *Bacteroides* colonization in both **(A)** serum and **(B)** lung tissue. The volcano plot shows the $\log_2$ fold change in cytokine expression between nonCF mice gavaged with *Bacteroides* versus PBS, plotted against the negative $\log_{10}$ transformed P-value of the difference in cytokine expression. Each point represents a cytokine, with the color indicating the direction and significance of the expression change. The legend corresponds to both panels. Cytokines with a significant increase in level ($P < 0.05$) in mice gavaged with *Bacteroides* compared to PBS are colored in blue, while cytokines with a significant decrease in level are colored in red. Cytokines with a p-value $< 0.05$ are colored in gray. Line representations include a $\pm$ 0.6 $\log_2$ fold change threshold (gray dashed vertical lines) and a p-value threshold of 0.05 (gray dashed horizontal line). The cytokine data were analyzed using a linear model for each cytokine, accounting for the effects of Treatment, Sex, their interaction and accounting for cage.

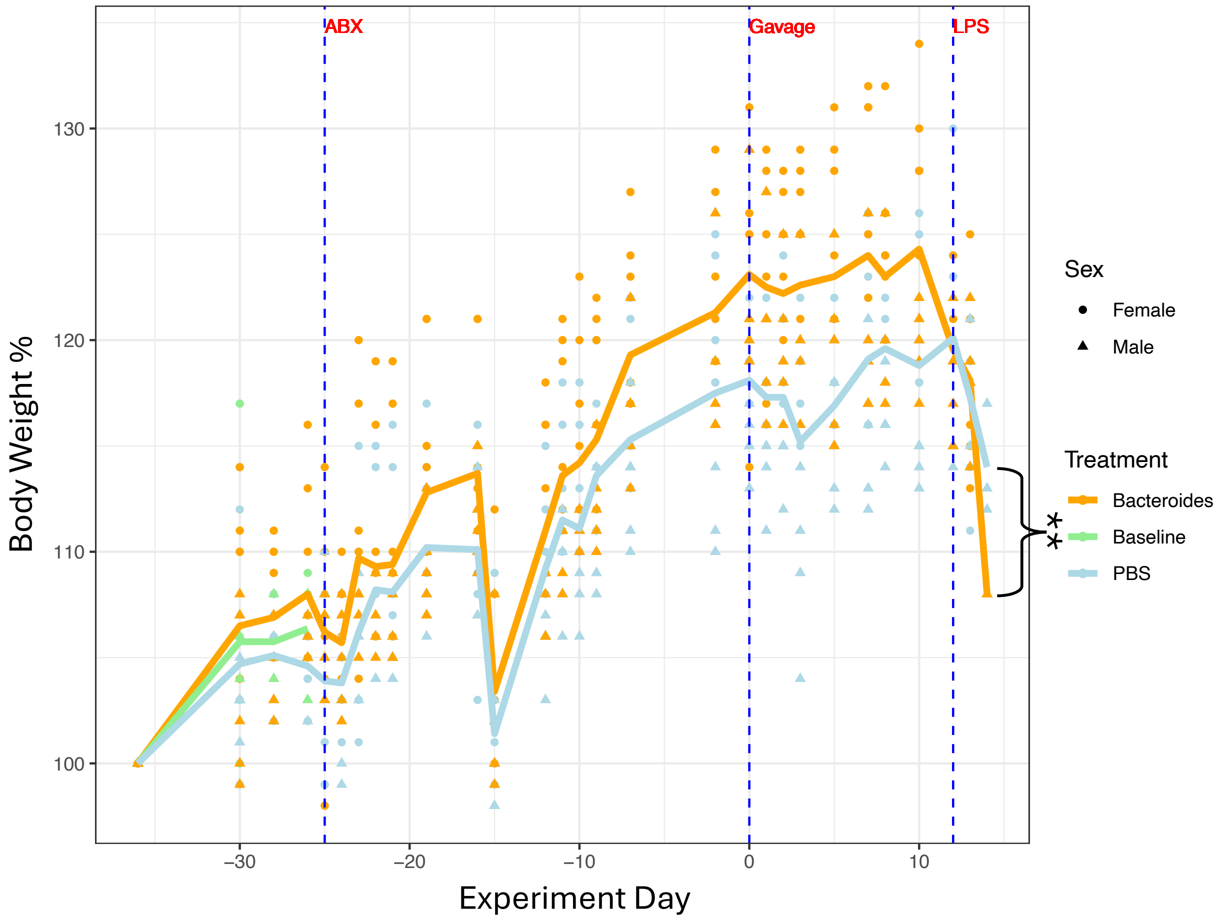


**Figure S9. *Bacteroides*-gavaged mice experience more weight gain.** Mouse body weight was recorded throughout the experiment. Weight is expressed as a percentage relative to each mouse's baseline weight at the start of the experiment. Each point represents an individual mouse, with sex indicated by point shape. The bold line denotes the average weight per treatment group over time. Statistical significance was assessed using a mixed effects model of weight percentage across treatment periods (Baseline, Antibiotics, Gavage), accounting for Treatment, Sex, their interaction, cohousing, and repeated measures. Following *Bacteroides* gavage, mice gained significantly more weight compared to PBS-treated mice ($p$ = 0.00495). Significance code: **, P < 0.01.

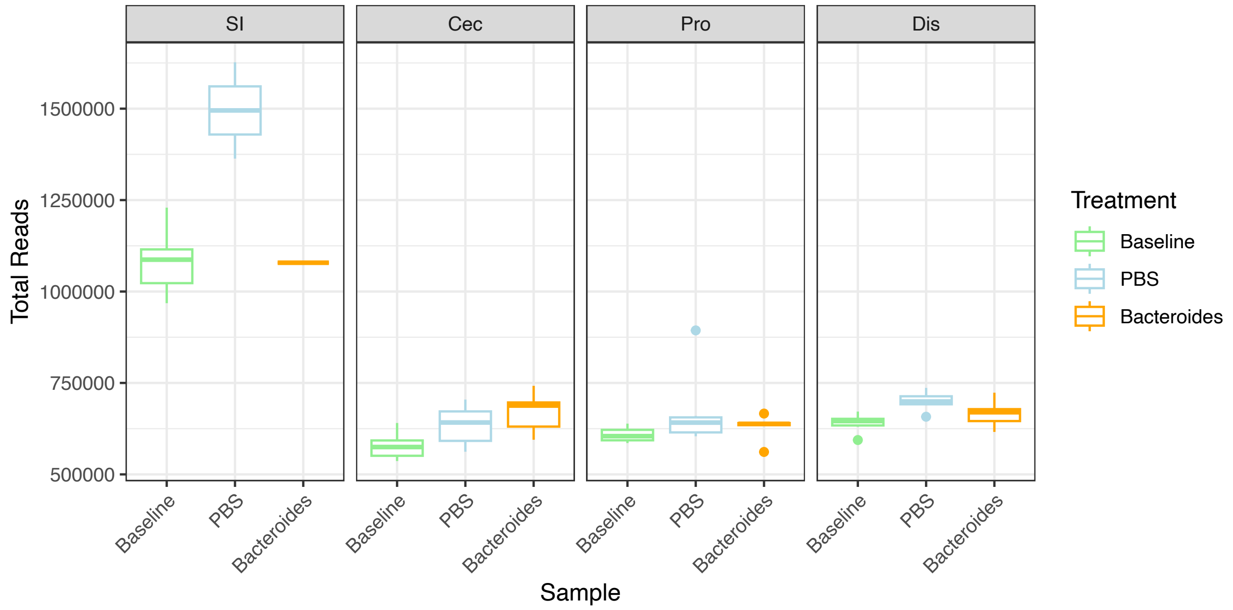


**Figure S10. NanoString counts in murine intestinal regions by treatment group**. This boxplot displays the total normalized NanoString counts of microbial RNA across different biogeographical regions of the mouse intestine—small intestine (SI), cecum (Cec), proximal colon (Pro), and distal colon (Dis)—stratified by treatment groups (Baseline, PBS, *Bacteroides*). The color coding of the boxplots corresponds to the treatment type, providing a visual comparison of total NanoString counts across the various intestinal segments.
