## Supplemental Table S2 for "Sex and regional effects of *Bacteroides* in the gut"

### Table S2: Number of genes differentially expressed by *Bacteroides* across the gut[a].

| | Small Intestine | Cecum | Proximal Colon | Distal Colon |
|---|---|---|---|---|
| Treatment | 60 | 55 | 23 | 26 |
| Sex | N/A | 122 | 23 | 13 |
| Combined Effect | N/A | 31 | 0 | 10 |
| Top Pathways Regulated | Chemokine Signaling, TCR Signaling, NK-kappaB Signaling | Myeloid Activation, NLR Signaling, Coagulation | NK Activity, Oxidative Stress Response, RNA Sensing | Myeloid Activation, Chemokine Signaling, BCR Signaling |

[a]For each region, genes significantly differentially expressed in each of the treatment groups of mice gavaged with PBS or Bacteroides were identified. Significantly differentially regulated genes were determined by performing a mixed-effects model on the $log_2$ transformed counts of each gene within each region, accounting for treatment, sex, and the combined effect of sex and treatment. Total number of genes found to be significantly affected by treatment, sex, and the combined effect of treatment and sex are listed in this table. Additionally, pathways that describe the function of each region's top 3 differentially expressed genes are highlighted in the last row.
